# High-throughput functional characterization of enhancers in totipotent-like cells

**DOI:** 10.1101/2025.11.03.686172

**Authors:** Lingyue Yang, Tianran Peng, Yating Zhu, Xuzhao Zhai, Boyan Huang, Ling Li, Tao Zhang, Jiekai Chen, Dan Liang, Jiangping He, Man Zhang

## Abstract

Zygotic genome activation (ZGA) marks the initial transcription event in embryogenesis, yet the *cis*-regulatory mechanism remains unclear. Here, utilizing massively parallel reporter assays, we functionally dissect enhancers across mouse genome in DUX-induced 2C-like cells (2CLCs). Integrated analysis with epigenomic and transcriptomic data from 2CLCs and 2-cell (2C) embryos, active enhancers in totipotent cells are depicted. Among them, a notable proportion of promoters exhibit enhancer activities, showing elevated active chromatin features and correlating with enhanced gene expression during ZGA. Furthermore, only half of the MT2_Mm exhibit enhancer activities in 2CLCs. In addition, 2CLC enhancers augment transcription in 2C embryos. Finally, deleting enhancer regions in both 2CLCs and 2C embryos highlighted their crucial role in facilitating transcription of ZGA genes. These findings advance our comprehension of the *cis*-regulatory mechanism governing ZGA process.

## Introduction

During embryogenesis, the zygote is formed by the fusion of two specified gametes ^1^. The zygotic genome is initially transcriptionally quiescent, allowing the maternal components in the oocyte to reset the genome into a naïve and globally accessible state ^2^. Subsequently, the zygotic genome is activated to establish a so-called totipotent state, enabling the resultant blastomeres with the potential to differentiate into all cell types of an organism, through a process known as the zygotic genome activation (ZGA) ^3^. ZGA is a critical developmental milestone as it signifies the beginning of embryonic genome exerting control over its own development ^4,5^, analogous to the initiation of a precisely programmed genetic code. Elucidating the mechanisms by which specific genes are selected for activation during ZGA is fundamental to understand the initiation of life. Despite its importance, the regulatory mechanisms governing mammalian ZGA remain poorly understood due to the limited materials.

Interestingly, it was reported that a small population of cells within mouse naïve embryonic stem cells (ESCs) show the activity of MERVL (murine endogenous retrovirus with leucine tRNA primer) elements and share hundreds of transcripts that are specifically activated in 2-cell (2C) embryos. These cells are therefore referred to as 2C-like cells (2CLCs) ^6–8^. Moreover, 2CLCs also exhibit chromatin architecture resembling that of the 2C embryos ^9–11^. These 2CLCs provide an opportunity for addressing the mechanism of ZGA. With the aid of this *in vitro* pluripotent-to-totipotent transition model, several critical transcriptional factors (TFs) and signaling ^12^ function in mammalian ZGA have been identified ^9,13–15^. Recently, several works reported that high proportion of 2CLCs can be generated from pluripotent cells by either ectopic overexpression of TFs ^16^ or culturing in a certain medium which contained epigenetic modifiers or the agonist of retinoic acid (RA) signaling ^17–20^. These systems expand the application of 2CLCs to study the mechanism underlying ZGA.

Enhancers, as vital *cis*-regulatory elements, are fundamental in orchestrating the spatially and temporally precise regulation of gene expression programs during embryonic development ^21^. In recent years, with the rapid development of low-input, high-throughput sequencing technology, the epigenetic modification profiles of early embryos are unveiled to predict the *cis*-regulatory modules ^22,23^. Nonetheless, the prediction based on epigenetic features may lead to false positive candidates. Putative enhancers need to be functionally validated through direct transcriptional reporter assays. Self-transcribing active regulatory region sequencing (STARR-seq) is a newly-developed massively parallel reporter assay which is able to quantitatively identify transcriptional enhancers directly based on their activity in a big scale by cloning sheared DNA between a minimal-promoter and a downstream polyA sequence ^24,25^. If the DNA fragment exhibits enhancer activity, this will result in transcription of the enhancer sequence. In this work, we performed STARR-seq across the whole murine genome in DUX-induced 2CLCs. Combining with the epigenetic modifications and transcriptome analysis in cells and 2C embryos, we depicted the potential functional enhancers in 2CLCs and ZGA process.

## Results

### STARR-seq dissects functional enhancers across murine genome in DUX-induced 2CLCs

In order to obtain sufficient 2C-like cells to conduct the genome-wide enhancer reporter assay, we established an *in vitro* DUX-induced 2CLC model. An ESC line containing a doxycycline (Dox)-inducible *Dux* transgene and a MERVL-LTR (LTRs, long terminal repeats) driven EGFP-F2A-puroR cassette (an indicator of the 2C-like state) was generated, which referred as iDux-MGP cells (Supplementary Fig. 1a). Puromycin was used at 6 hours after Dox supplementation to help eliminate cells in which MERVL activation did not occur (Supplementary Fig. 1a). Single cell derived clones from iDux-MGP cells were then picked up, and clone 3 which exhibited high 2CLC induction efficiency and normal karyotype was used for the following experiments (Supplementary Fig. 1b, c). To induce 2CLCs, doxycycline was added for 30 hours (Supplementary Fig. 1a, b). Transcriptome analysis including mRNA and long non-coding RNA (lncRNA) in sorted DUX-induced 2C::EGFP-positive (GFP^+^) and -negative (GFP^−^) cells identified 4,994 transcripts upregulated in GFP^+^ cells, including 2C specific markers such as *MERVL-int*, *MT2_Mm*, *Zfp352*, and *Zscan4* family (Supplementary Fig. 1d, e and Supplementary Table 1).

To further assess the properties of cells with or without DUX induction in a single-cell level, we performed single-cell RNA-seq (scRNA-seq) in the iDux-MGP-Clone 3 cells with or without Dox treatment. By integrating our scRNA-seq data with in-house dataset of mouse early embryos^26^ and other published datasets from totipotent cell types, including reported DUX-induced 2C-like cells (2CLCs), totipotent-blastomere like cells (TBLCs) and chemically induced totipotent stem cells (ciTotiSCs) ^16,19,20^, we found that both DUX-induced 2CLCs and ciTotiSCs were mainly clustered with the cells of 2-4 cell (2C-4C) embryos, with a subset of DUX-induced 2CLCs close to an intermediate population reported previously (Supplementary Fig. 1f). In contrast, the -Dox cells mainly centered near epiblast and mESCs (Supplementary Fig. 1f). Furthermore, unsupervised clustering analysis identified six distinct clusters. It showed that 59.7% of the -Dox group cells were in Cluster 1, where epiblast and mESCs were located, and 22.66% of cells were in Cluster 4, which closed to the inner cell mass (ICM) cells (Supplementary Fig. 1g). Only 4.51% of the -Dox cells were in Cluster 2, where the 2C-4C embryos and reported 2CLCs located. Conversely, 59.33% of the +Dox cells were located in Cluster 2, while only 4.46% were in Cluster 1 and 3.77% in Cluster 4 (Supplementary Fig. 1g). The gene expression UMAP plot indicated that GFP^+^ cells mainly located in Cluster 2 and enriched 2C markers such as *Zscan4c*, *Zscan4d*, and *Ddit4l* (Supplementary Fig. 1h). These data demonstrate that our induction system effectively generated a high proportion of 2CLCs, which can be used for the further studies.

To identify potential functional enhancers across the whole murine genome in totipotent-like cells, we constructed a STARR-seq plasmid library by inserting sonication-sheared genomic DNA at the average size of 700-1200 bp into the optimized STARR-seq vector ^27^. The plasmid library was then transfected into the cells at 24 hours after the Dox addition and cells were collected at 16 hours after the transfection (Fig. 1a). The plasmids and the mRNAs transcribed from the library were isolated respectively from the transfected cells for sequencing, and the plasmids were set up as the input (Fig. 1a). Two independent experiments were performed. Input DNA coverage analysis revealed adequate representation of the mouse genome in our library, with one replicate surpassing 4 times of the genome coverage and the other exceeding 11 times (Supplementary Fig. 2a). Moreover, Pearson’s correlation analysis indicated strong reproducibility between the two replicates of the STARR-seq data (Supplementary Fig. 2b). Fragment sequences which exhibited self-transcription levels at least 2-fold higher than those of the input were defined as STARR peaks, which were significantly enriched over the input compared to the randomly selected fragments from genome (Supplementary Fig. 2c). The sizes of STARR peaks corresponded with those of the input, predominantly falling within the range of 700-1200 bp, without interference from smaller fragment noise (Supplementary Fig. 2d). In addition, we generated ATAC-seq and ChIP-seq libraries (targeting H3K4me1, H3K4me3, and H3K27ac) in sorted GFP^+^ and GFP^−^cells following DUX induction, aiming to combine the analysis of STARR peaks with the chromatin status and epigenetic modifications in 2CLCs (Supplementary Fig. 2e, f).

**Fig. 1.**
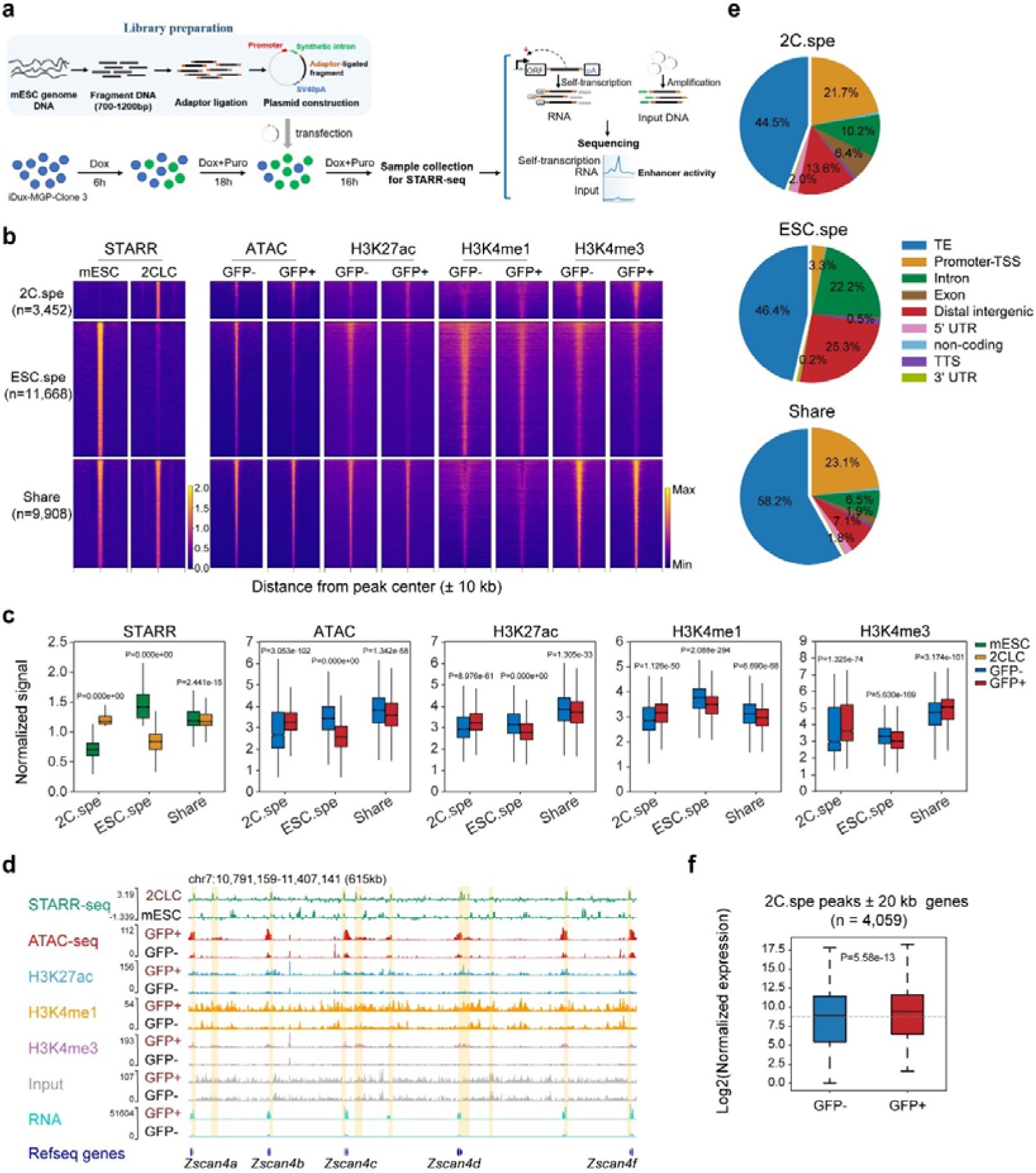
STARR-seq dissects functional enhancers across murine genome in 2CLCs. **a.** Schematic of whole-genome STARR-seq in DUX-induced 2C::EGFP positive cells. **b.** Heatmap showing the differential STARR peaks between 2CLCs and mESCs, and combined with indicated multi-omics analysis of chromatin characteristics in 2C::EGFP negative (GFP-) and 2C::EGFP positive (GFP+) cells (see Supplementary Fig. 2e). Signal was computed on the STARR-seq peaks flanked by 10 kb. 2C.spe: specific STARR peaks in 2CLCs; ESC.spe: specific STARR peaks in mESCs; Share: STARR-peaks in both 2CLCs and ESCs. **c.** Boxplot showing the overall signal intensity of indicated marks in Fig. 1b in indicated cell types. Boxplots denote the medians and the interquartile ranges (IQR). The whiskers of a boxplot are the lowest datum still within 1.5 IQR of the lower quartile and the highest datum still within 1.5 IQR of the upper quartile. P-values were from a Mann-Whitney U test. **d.** A genomic snapshot showing STARR-seq, ATAC-seq, ChIP-seq of the specified histone modification with input, and RNA-seq tracks at the loci of *Zscan4* family in indicated cells. **e.** Pie charts showing the genomic distribution of the STARR-peaks in Fig. 1b. **f.** Boxplot showing the expression levels of the genes flanked by 2C-specific (2C.spe) STARR peaks within 20 kb (± 20kb) in 2C::EGFP negative (GFP-) and 2C::EGFP positive (GFP+) cells. P-values were from a Mann-Whitney U test.

To identify the 2C-specific enhancers, we conducted an integrative analysis of enhancers in 2CLCs and mESCs using a published STARR-seq dataset of mESCs (GSE143546) ^28^. Given that enhancers detected by transient-assay are based on exogenous plasmids, and chromatin is a complex structure, highly active enhancers typically reside in open and active chromatin regions to effectively regulate target genes ^29^. Therefore, we firstly classified enhancers based on their STARR activities and chromatin opening status. Among them, putative enhancers exhibiting STARR activities and locating in open chromatin regions in either 2CLCs or mESCs were defined as Cluster 1 (C1). In contrast, elements with STARR activities yet residing in close chromatin regions in the respective cell types were defined as C2, suggesting that their enhancer activities probably be suppressed in cells. In addition, elements located in active chromatin regions without STARR activities were defined as C3, implying that these regions although had the active epigenetic modifications, might not act as enhancers in cells (Supplementary Fig. 3a). Therefore, our subsequent functional enhancer analysis in 2CLCs focused primarily on the STARR peaks in C1. Subsequently, the differential STARR peaks were calculated between the 2CLCs and mESCs in cluster C1 (n = 25,028). Using a 2-fold threshold, we identified 3,452 2C-specific enhancers (2C.spe), 11,668 ESC-specific enhancers (ESC.spe) and 9,908 shared enhancers in 2CLCs and mESCs (Share) (Fig. 1b and Supplementary Fig. 3b, c). Integrated analysis with epigenomic data indicated that 2C-specific enhancers exhibited elevated signals of ATAC, H3K27ac, H3K4me1 and H3K4me3 in DUX-induced GFP^+^ cells, such as the STARR peaks within the *Zscan4* family gene loci (Fig. 1b-d). Whereas, ESC-specific enhancers were associated with increased signals in GFP^−^cells (Fig. 1b, c). These results suggested that the enhancer sequences identified via STARR-seq align with the chromatin features of enhancers in both cell types. Genome annotation revealed a significant enrichment of transposable element (TE) components in the three groups of enhancers (Fig. 1e). Notably, over 20% of enhancers in 2CLCs were located within promoter regions, while only 3.3% of ESC-specific enhancers were situated in promoter regions (Fig. 1e). Moreover, among the genes expressed in DUX-induced GFP^+^ cells, those promoter regions with enhancer activities had higher levels of epigenetic modification related to active chromatin features, compared to those without enhancer activities (Supplementary Fig. 3d). Furthermore, we found that the expression levels of genes around 2C-specific enhancers (± 20kb) were significantly higher in DUX-induced GFP^+^ cells (Fig. 1f), especially for non-coding RNAs (Supplementary Fig. 3e), suggesting their functional role in regulation of gene expression in 2CLCs.

Enhancers encompass the binding sites to recruit cell-type specific transcription factors (TFs) for facilitating the initiation of transcription ^30^. We then conducted a motif analysis within each class of C1 enhancers to examine the TFs potentially involved in these interactions. It revealed that ESC-specific enhancers were enriched for pluripotency TFs, such as OCT4, SOX2, and members of KLF family. Shared enhancers were enriched some pan-TFs, such as PRDM14 and YY1. Whereas, 2C-specific enhancers were particularly enriched the TFs highly expressed during the ZGA process, including TEAD4, DUX and GATA2, with the latter two playing vital roles in activating ZGA genes in previous studies ^9,13,31,32^ (Supplementary Fig. 3f, g). These findings underscore the distinct characteristics between 2C-specific and ESC-specific enhancers. Taken together, we identified in total 13,360 enhancer elements in 2CLCs by using STARR-seq in conjunction with multi-omics analysis, and further examined the distinct characteristics between 2C-specific and ESC-specific enhancers.

### Evaluation of the *in vivo* functional potential of enhancers identified in 2CLCs

To investigate the *in vivo* functional potential of the enhancers we identified in 2CLCs, we first examined the expression profiles of genes associated to the 2C-specific (2C.spe) enhancers (± 20kb) in embryos, using previously published transcriptome data from different stages of embryos ^2^. The results showed that the associated genes of 2C-specific (2C.spe) enhancers exhibited a significant increase in expression at early 2C (E2C) stage, followed by a striking decrease in expression in ICM (Supplementary Fig. 4a). Subsequently, we conducted an integrated analysis of the three classes of C1 enhancers with the published ATAC-seq data in early embryos ^2^ (Fig. 2a). Interestingly, consistent with their differential activities *in vitro*, enhancers identified in 2CLCs (2C.spe and share) had higher levels of transposase-accessibility in the E2C and 2C stages compared to the ESC-specific enhancers. Whereas, the ATAC-seq signals associated with ESC-specific enhancers were much lower compared to the enhancers identified in 2CLCs in E2C and 2C stage. Interestingly, while the signals of 2C-specific enhancers progressively decreased to their lowest point in ICM after they reached their culmination at 2C stage, the ATAC-seq signals associated with ESC-specific enhancers dramatically increased at 8-Cell stage and maintained their high levels in the ICM of embryos (Fig. 2a). Specifically, 42% of 2C-specific enhancers and 22% of shared enhancers were in an open status in 2C embryos, while only 17% of ESC-specific enhancers were located in the open chromatin and with a lowest level of ATAC-seq signal in 2C embryos (Supplementary Fig. 4b, c). These results suggest the *in vivo* functional potential of the enhancers identified in 2CLCs.

**Fig. 2.**
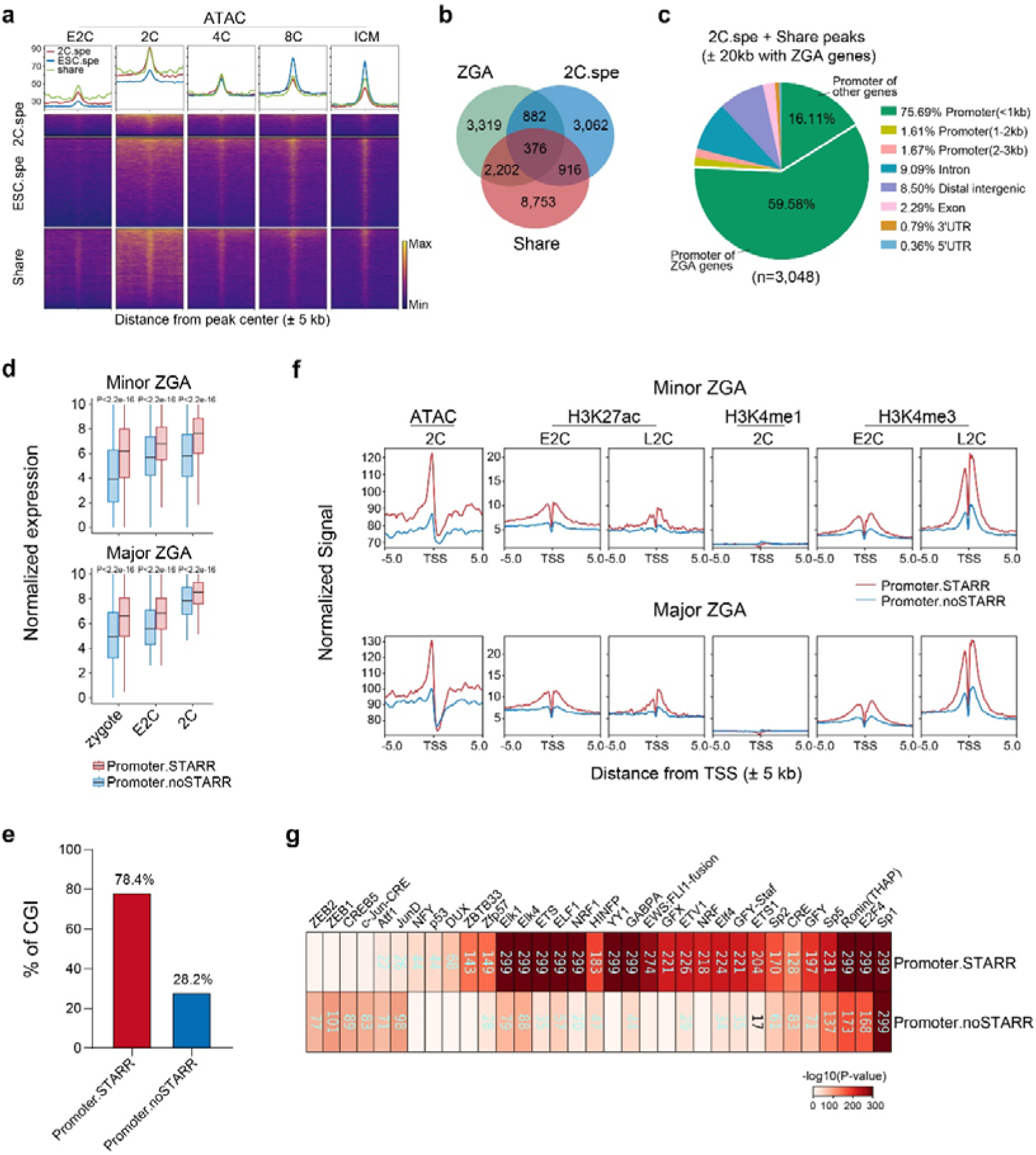
Evaluation of the *in vivo* functional potential of enhancers identified in 2CLCs. **a.** Heatmap showing the ATAC-seq signals of the indicated groups of STARR peaks during mouse early embryonic development. Signal was computed on the STARR-seq peaks flanked by 5 kb. The ATAC-seq data were obtained from published data (GSE66390). **b.** Venn diagram illustrating the overlap between ZGA genes and genes flanked by 2C-specific (2C.spe) or shared (Share) STARR peaks within 20kb (± 20kb). **c.** Pie charts illustrating the genomic distribution of 2C-specific (2C.spe) and shared (Share) STARR peaks in proximity to ZGA genes, defined with STARR peaks located within 20 kb (± 20kb) of ZGA gene loci. **d.** The expression levels of minor (top) and major (bottom) ZGA genes categorized by the presence (Promoter.STARR) or absence (Promoter.noSTARR) of STARR-seq signals at their promoter regions in 2CLCs. The number of minor and major ZGA genes with STARR-seq signals at the promoter regions (Promoter.STARR) is 1,116 and 1,392 respectively. **e.** The CpG island distribution of the promoter sequences (Promoter.STARR and Promoter.noSTARR). Promoter.STARR: the ZGA gene promoters with STARR-seq signals in 2CLCs; Promoter.noSTARR: the ZGA gene promoters without STARR-seq signals in 2CLCs. **f.** Line chart showing ATAC-seq, H3K27ac, H3K4me1 and H3K4me3 signals at the indicated promoter regions (TSS ± 5 kb) of minor (top) or major ZGA genes (bottom) in 2C embryos, categorized by the presence (Promoter.STARR) or absence (Promoter.noSTARR) of STARR-seq signals in 2CLCs. E2C: early 2-cell embryos; L2C: late 2-cell embryos. **g.** Heatmap showing transcription factor (TF) motifs identified from ZGA gene promoters (TSS ± 1kb), categorized by the presence (Promoter.STARR) or absence (Promoter.noSTARR) of STARR-seq signals in 2CLCs. The color indicated the enrichment of the motifs (-log10 P-value).

To further investigate potential target genes of the identified 2C enhancers in 2C embryos, we first defined 4,744 minor and 3,446 major ZGA genes by analyzing differentially expressed genes between E2C/Zygotes and MII oocytes as well as between late 2C (L2C) and early 2C (E2C) embryos respectively with previously published data ^2^ (Supplementary Fig. 4d, e and Supplementary Table 2). In total, 3,460 ZGA genes were associated to the enhancers identified in 2CLCs (2C.spe and share), up to 51% (3,460/6,779) of the total ZGA genes (Fig. 2b). Notably, genome annotation of the STARR-peaks in 2CLCs around ZGA genes (± 20 kb) revealed that 75.69% of the peaks were located in the gene promoter regions, with 59.58% of these genes being ZGA genes (Fig. 2c and Supplementary Table 3). Given that the 3D chromatin structure has not been fully established in mouse 2C embryos ^33,34^, which results in the weakening regulation of distal enhancers, especially in minor ZGA ^35^, those enhancer-promoters might be an alternative way in early embryos to promote gene transcription. Indeed, genes with enhancer-promoters had a significant higher expression level compared to those whose promoters had no enhancer activities (Fig. 2d). Previous works suggested that promoters of some housekeeping (HK) genes exhibit enhancer activities in flies and pluripotent stem cells (PSCs) ^25,36^. Nevertheless, with a reference of HK genes in mouse ^37^, our data revealed that 60.18% promoters of ZGA genes are non-HK genes. Notably, these genes included the critical TFs during ZGA process, such as *Dux*, *Ddit4l*, *Zfp352* and *Zscan4* family (Supplementary Fig. 4f and Supplementary Table 3). Moreover, GO analysis revealed that the non-HK ZGA genes with enhancer-promoters predominantly enriched in ribosome biogenesis and nuclear division, which were essential for cell survival and important for maternal-to-zygotic transition (MZT) (Supplementary Fig. 4g).

To investigate what distinguishes the enhancer-promoters from other promoters of ZGA genes, we first characterized their sequences. While both classes of promoters exhibited similar level of GC content (Supplementary Fig. 4h), 78% enhancer-promoters contained CpG islands (CGI), in contrast only 28% of promoters without STARR activities had CGI (Fig. 2e). Subsequently, we compared the chromatin status ^2^ and epigenetic modifications ^22,23^ between these two different groups of promoters. The results revealed that promoter regions with enhancer activities of both minor or major ZGA genes exhibited much higher levels of ATAC-seq signal in 2C embryos compared to promoters without STARR activities (Fig. 2f). Moreover, the enhancer-promoter regions displayed much higher levels of H3K27ac and H3K4me3 in E2C and L2C embryos than those without STARR peaks (Fig. 2f). Nevertheless, as the global H3K4me1 levels were weak in 2C embryos (Supplementary Fig. 4i), the levels of H3K4me1 were low in both promoters of ZGA genes in 2C embryos (Fig. 2f). To further explore the potential mechanism underlying the enhancer-promoters, a motif analysis was performed. Interestingly, Additional TFs exclusively enriched in the enhancer-promoters were identified, including NRF1 and NFY, the two well-known transcription activators ^38–40^, as well as GABPA, P53 and DUX4, the key regulators of ZGA genes ^41,42^ (Fig. 2g). Notably, most of these TFs specifically highly expressed in MII oocytes or during ZGA process (Supplementary Fig. 4j).

### MERVL elements with enhancer activities were dissected in totipotent cells

It was reported that transposable elements (TEs), especially MERVL elements, play an important *cis*-regulatory function in the regulation of ZGA genes, and therefore, are critical for embryonic development ^6–8,43–45^.To investigate the enhancer function of TEs, we first annotated the TE families within the 2C-specific and ESC-specific enhancers. The results showed that the ERVL (ERV3) family, such as MT2_Mm, MT2C_Mm and ORR1A1-int, were predominantly enriched in the 2C-specific enhancers (Fig. 3a-c). While ESC-specific enhancers were primarily enriched elements of ERVK and ERV1, like ETnERV, MMETn-int and LTRIS2 (Fig. 3a, b and Supplementary Table 4). These results indicated different TE clusters were adopted as enhancer elements in these two different cell types.

**Fig. 3.**
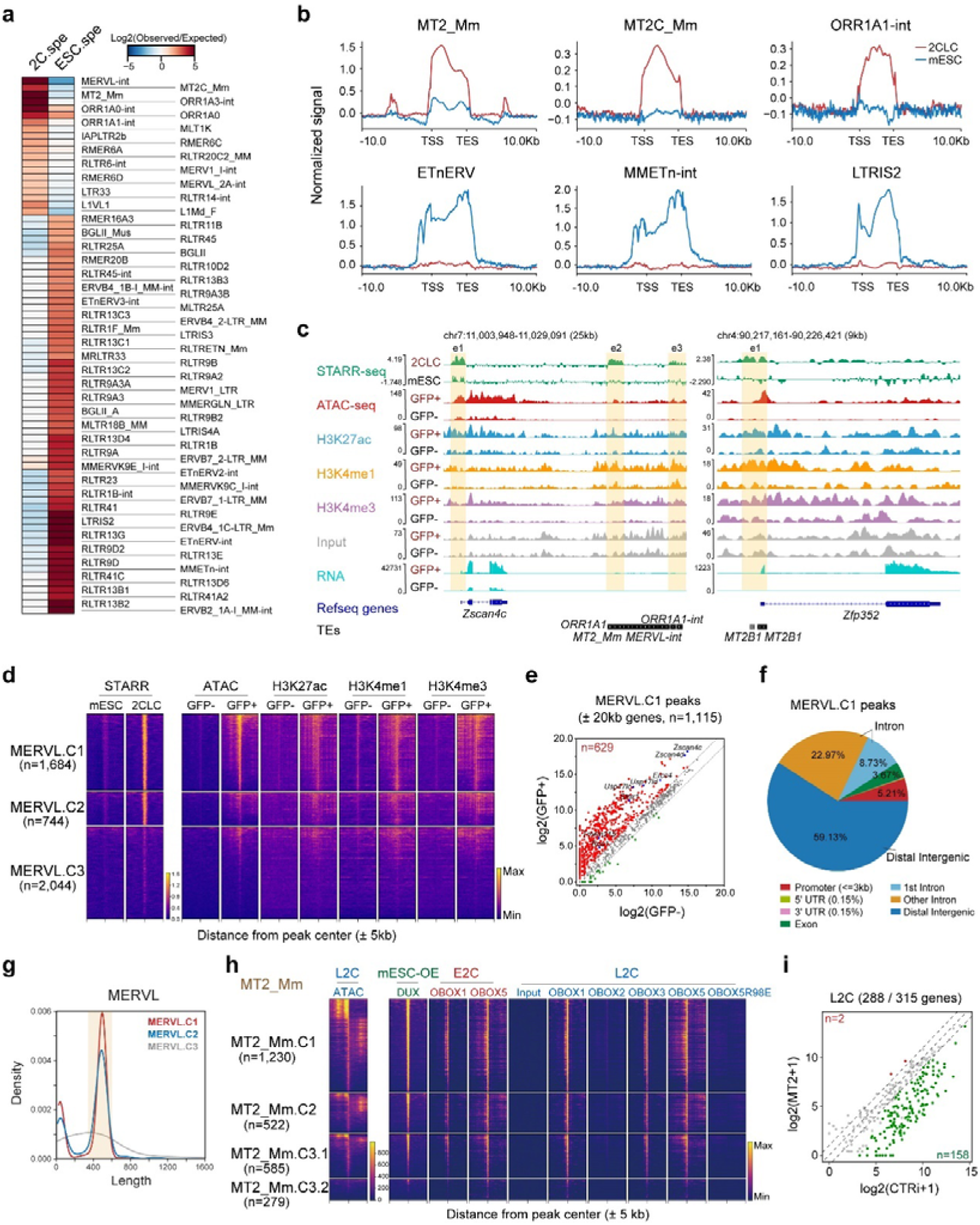
MERVL elements with enhancer activities were dissected in totipotent cells. **a.** Heatmap showing the enrichment of TEs in the 2C-specific (2C.spe) and ESC-specific (ESC.spe) STARR peaks in Fig. 1b. The fold enrichment was calculated by the ratio of the observed over expected counts of the modified TEs among randomly selected genomic regions. **b.** Read count tag density pileups (in RPKM) of the STARR-seq signals for the selected TEs in 2CLCs and mESCs. The TEs were scaled to the same size, and the flanking 10 kb regions are shown. **c.** Genomic snapshots showing STARR-seq, ATAC-seq, ChIP-seq of the specified histone modification with input, and RNA-seq tracks at the indicated regions with genes and TEs in the indicated cells. The 2C-specific enhancers are highlighted (Yellow square). **d.** Heatmap illustrating the classification of all MERVL elements into three clusters based on STARR-seq and ATAC-seq signals in 2C-like cells (2CLCs) (left). MERVL.C1: cluster comprises MERVL elements showing both high STARR-seq signals and ATAC-seq signals in MERVL-GFP^+^ cells. MERVL.C2: cluster includes MERVL elements with STARR-seq signals and minimal ATAC-seq signals in MERVL-GFP^−^cells. MERVL.C3: cluster consists of MERVL elements exhibiting low or no STARR-seq signals in 2CLCs. The right panel displays the indicated histone modifications associated with these three clusters of MERVL elements. **e.** Scatter plot showing the expression levels of genes flanked by the MERVL.C1 elements in Fig. 3d within 20 kb (± 20kb). The X-axis represents the expression levels in GFP^−^ cells, while the Y-axis denotes the expression levels in MERVL-GFP^+^ cells. Each point represents a gene. Red dots highlight the upregulated genes in MERVL-GFP^+^ cells. **f.** Pie chart showing the genome distribution of the MERVL.C1 elements in Fig. 3d. **g.** Length distribution of the three clusters of MERVL elements in Fig. 3d. **h.** Heatmap showing MT2_Mm from the corresponding MERVL clusters (as shown in Fig. 3d) were classified into four clusters based on their ATAC-seq signals in L2C embryos (left). The binding of indicated transcription factors (TFs) in these four clusters of MT2_Mm elements were also shown (right). **i.** The differential expressed genes between MT2_Mm inhibited late 2-Cell (L2C) embryos and control embryos within the MERVL.C1 associated ZGA genes (n = 315) in Supplementary Fig. 5i. MT2_Mm were perturbed by injection of dCas9-KRAB with six MT2_Mm-targeting gRNAs. Control embryos were injected dCas9-KRAB with non-targeting gRNAs. Green dots highlight the downregulated genes in 2C embryos with MT2_Mm perturbation. The RNA-seq data from previous work were used (GSE242121).

Considering that there are nearly 4,472 sequence-conserved copies of MERVL and their LTRs within the mouse genome according to the RepeatMasker, the locations and characteristics of the MERVL elements with enhancer activities are still enigmatic. To this end, we further dissected the MERVL elements including MERVL internal regions (MERVL-int), and their LTRs (MT2_Mm) in our STARR peaks. Although the sequences of MERVL elements are highly repetitive, we found that not all MERVL repeats had enhancer activities (Supplementary Fig. 5a). Consequently, we divided the MERVL elements into three clusters based on their STARR activities and chromatin openness in 2CLCs (Fig. 3d). Cluster 1 MERVL elements (MERVL.C1, n = 1,684) exhibited high STARR activities, and co-localized with the transposase-accessible regions in 2CLCs, which were considered as functional enhancers in 2CLCs. Cluster 2 (MERVL.C2, n = 744) represented MERVL repeats with STARR-seq signals but low chromatin openness in 2CLCs, suggesting that although these repeats had enhancer potential, they were unlikely able to function as active enhancers in 2CLCs. Cluster 3 (MERVL.C3, n = 2,044) represented MERVL repeats with nearly undetectable STARR-seq signals in 2CLCs, and were unlikely to function as enhancers (Fig. 3d). Consistent with their function in 2CLCs, MERVL.C1 had much higher levels of chromatin openness and active histone modifications, including H3K27ac, H3K4me1 and H3K4me3, in GFP^+^ cells compared to those of MERVL.C3 (Fig. 3d and Supplementary Fig. 5b). Moreover, 56% (629/1,115) of MERVL.C1 associated genes (± 20 kb) were upregulated in DUX-induced GFP^+^ cells, including *Zscan4* and *Usp17* families (Fig. 3e and Supplementary Table 5).

To characterize the MERVL elements with enhancer activities, genome annotation revealed that majority of them were situated in distal intergenic regions (59%) and introns (31.6%), with 5.2% of them locating in annotated gene promoters (Fig. 3f), suggesting their distal regulatory function. Size distribution analysis revealed that except a small fraction of elements measuring nearly 100 base pairs (bp), majority of MERVL.C1 and C2 had an average size of around 500 bp, a size which is similar to the MERVL LTRs (MT2_Mm), suggesting MT2_Mm represents the majority of MERVL repeats with enhancer activities (Fig. 3g). Indeed, MT2_Mm represented 73.04% (1,230/1,684) and 70.16% (522/744) of the MERVL.C1 and MERVL.C2 respectively, whereas 42.27% in MERVL.C3 (Supplementary Fig. 5c). Further analysis revealed that the truncated MERVL-int (∼ 100bp) in MERVL.C1 were predominantly located near the LTRs with STARR activities, with 89.1% nearby LTRs in MERVL.C1 and 8.9% in MERVL.C2 (Supplementary Fig. 5d, e). These suggest that the detection of their enhancer activities is attributed to their spatial association with MT2_Mm. In addition, MT2_Mm in MERVL.C1 and MERVL.C2 consisted of either solo-LTR or the 5’ and 3’ LTRs of full length MERVL with no obviously biased. Whereas, MERVL.C3 had more solo LTRs (Supplementary Fig. 5f). As our analysis identified MT2_Mm as the primary elements of MERVL exhibiting enhancer functions, to investigate the enhancer potential of MERVL *in vivo*, we performed integrated analysis of MT2_Mm in the three MERVL clusters with published ATAC-seq in 2C embryos ^2^ (Fig. 3h). The results showed that almost all MT2-Mm in MERVL.C1 (designated as MT2_Mm.C1) had high chromatin openness in 2C embryos. Interestingly, despite MERVL.C2 exhibiting low ATAC-seq signal intensity *in vitro*, the majority of MT2_Mm within MERVL.C2 (designated as MT2_Mm.C2) displayed ATAC-seq signals in 2C embryos, a bit lower than those of MT2_Mm.C1, suggesting their potential functionality *in vivo*. In addition, albeit the absence of STARR activities of MERVL.C3 in 2CLCs, there were two distinct subpopulations of LTRs within MERVL.C3. Notably, 67.7% of these (designated as MT2_Mm.C3.1) showed chromatin accessibility in 2C embryos, while the other subpopulation (designated as MT2_Mm.C3.2) exhibited no chromatin accessibility *in vivo* (Fig. 3h and Supplementary Fig. 5g). Consistent with their chromatin accessibility *in vivo*, the LTRs which showed high intensity of ATAC-seq signals were also bounded by DUX in 2CLCs ^9^ and OBOX1/3/5 in 2C embryos ^46^. As a control, they were not bound by the OBOX2 and DNA-binding mutant OBOX5 (R98E) (Fig. 3h and Supplementary Fig. 5g). Given the highly conserved sequences of MT2_Mm (MERVL-LTRs), it is noteworthy to investigate the reasons why the LTRs within MT2_Mm.C3.2 were not bound by DUX and OBOX. Sequence alignment analysis revealed that while MT2_Mm with chromatin accessibility in 2C embryos were of full length, the MT2_Mm within MT2_Mm.C3.2 were truncated mutants, with most of them containing less than the first 150 bp of LTRs, lacking the DUX and OBOX binding motifs (Supplementary Fig. 5h). This might explain why this part of LTRs (MT2_Mm.C3.2) had no active enhancer function and low chromatin accessibility. Furthermore, 315 ZGA genes were associated to the MERVL.C1 (within ± 20 kb) (Supplementary Fig. 5i and Supplementary Table 5). Notably, half of them were downregulated upon the MT2_Mm perturbed by dCas9-KRAB with 6 small-guide RNAs (sgRNAs) ^44^, suggesting a direct regulatory influence of MT2_Mm on the expression of these genes (Fig. 3i and Supplementary Table 5).

### Verification of the 2C specific enhancers *in vitro* and *in vivo*

To further substantiate the function of the enhancers identified in 2CLCs *in vivo*, we performed an ultra-low-input STARR-seq (ULI-STARR-seq) in 2C embryos. This protocol had been specifically optimized from UMI-STARR-seq protocol ^47^ for low-input samples with an enhanced RNA recovery method. Due to the restricted availability of early embryos, it is unfeasible to evaluate all enhancers *in vivo.* Therefore, 88 candidates from the enhancers identified in 2CLCs (2C.spe or share), with their location proximal to ZGA-related genes such as *Zscan4f* (seq19) and *Usp12* (seq54) were synthesized and cloned into the STARR-seq plasmid (Fig. 4a and Supplementary Fig. 6a). Additionally, 9 DNA fragments without STARR and ATAC peaks in 2CLCs were cloned to serve as negative controls (Fig. 4a and Supplementary Fig. 6a). The constructed STARR-seq plasmid library containing both the candidate enhancers and negative controls, which was injected into the male pronucleus of zygotes, and self-transcriptional RNAs were then collected for sequencing at the L2C stage (Fig. 4a). Two independent experiments were conducted. It is noted that although activity was detected in two negative controls, it was not reproducible across replicates (Supplementary Table 6). The 7 remaining controls displaying no enhancer activities in both replicates were used for the following analysis. The results revealed that 69% of the candidate enhancers (61 out of 88) exhibited enhancer activities in both replicates, indicating their functional enhancer capacity in 2C embryos (Fig. 4b, c, Supplementary Fig. 6b and Supplementary Table 6). To cover the complexity of the fragments, in the library, the selected 88 candidates encompassed both TE and non-TE elements with diverse genome distribution (Supplementary Fig. 6c, d). Notably, a comparable proportion of LTRs, and enhancer-promoter candidates showed activities in 2C embryos, further validating their enhancer function *in vivo* (Supplementary Fig. 6e, f). To further validate the activities of the enhancers identified in the mini-STARR screening, fluorescence reporter plasmids were constructed. Control elements and 10 candidate enhancers with 4 promoter elements showing enhancer activities *in vivo* were inserted after the EGFP coding sequence in the reporter plasmid with a minimal promoter (Supplementary Fig. 6g). Each plasmid along with a constitutively expressed mCherry tracer plasmid were co-injected into the male pronucleus of zygotes. Fluorescence was subsequently observed and quantified at 24 hours after microinjection. The results confirmed that all the 10 candidate enhancers drove significantly higher GFP expression compared to the negative controls or the promoter elements showing no STARR activities in 2CLCs (promoters of *Duxbl1*, *Obox6*, *Klf4*, *Pou5f1*) (Supplementary Fig. 6h, i).

**Fig. 4.**
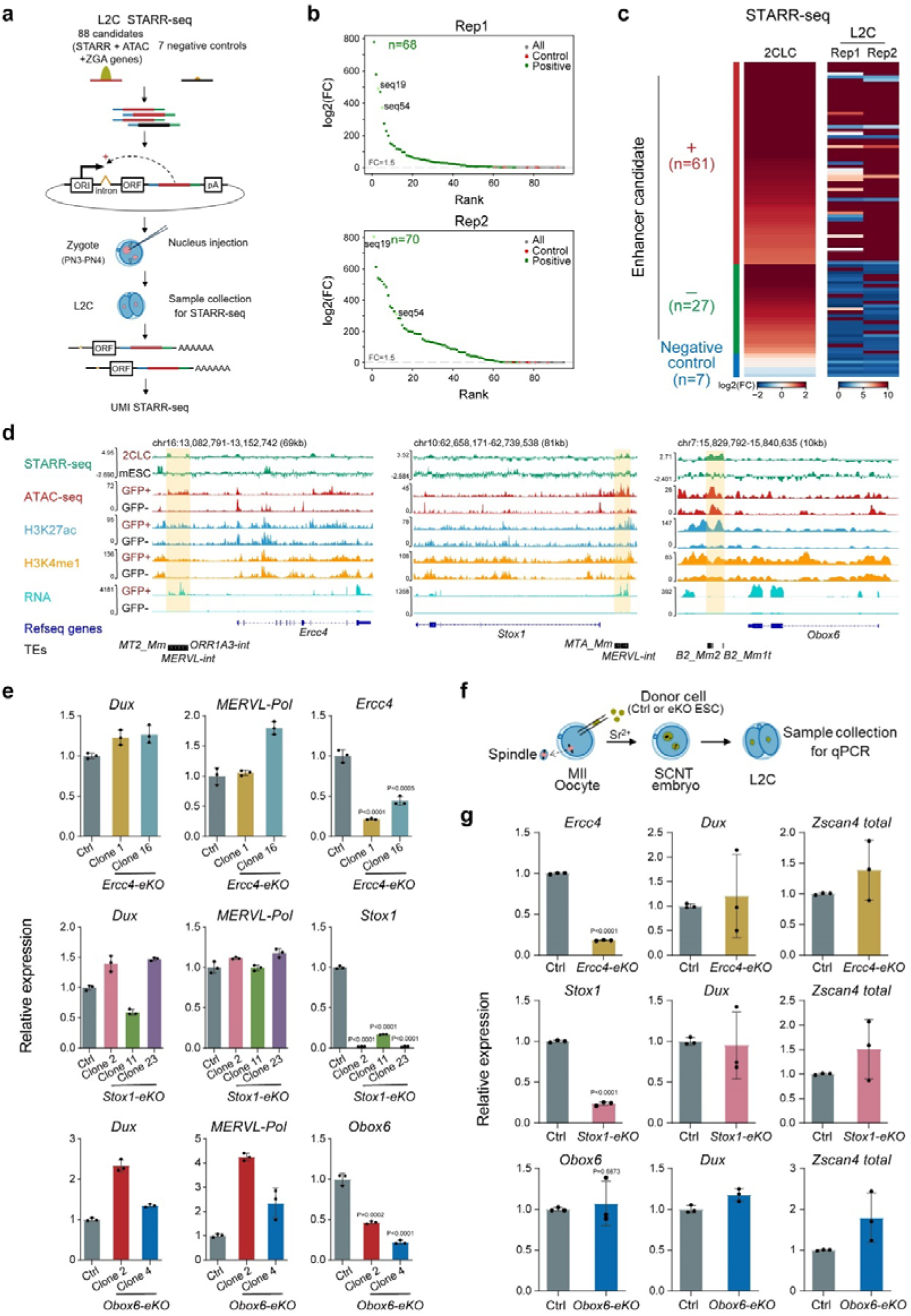
Verification of the 2C specific enhancers *in vitro* and *in vivo*. **a.** Schematic of ULI-STARR-seq in mouse L2C embryos. 88 candidates and 7 negative controls were manually cloned into the STARR-seq constructs. Pooled products were then injected into the male pronucleus of zygotes. L2C: late 2-cell embryos. **b.** Scatter plot representing the fold enrichment of enhancer candidates relative to the negative control in mouse L2C STARR-seq samples. The Y-axis shows the fold change divided by the median of negative control (red dots); values greater than 1.5-fold are considered positive enriched (green dots). The top panel shows data from replication 1 (Rep1) with 200 L2C embryos, and the bottom panel shows data from replication 2 (Rep2) with 480 L2C embryos. Enhancer candidates shown in Supplementary Fig. 6a (seq19 and seq54) were highlighted with light green. L2C: late 2-cell embryos. **c.** Heatmap displaying the intensity of STARR-seq signals for enhancer candidates (n = 88) and negative controls (n = 7) in 2C-like cells (2CLCs, left) and mouse L2C embryos (right). The heatmap visualizes the relative STARR-seq signal intensities across the samples, with rows representing each enhancer candidates and negative controls, and columns corresponding to different replicates in 2CLCs and L2C embryos. Enhancer candidates (n=61) that exhibited positive enrichment (+) in both replicates of the L2C samples are indicated, suggesting their potential enhancer activity in mouse L2C embryos. L2C: late 2-cell embryos. **d.** Genomic snapshots showing STARR-seq, ATAC-seq, ChIP-seq of the specified histone modification with input, and RNA-seq tracks at the indicated genome regions with the genes and TE elements labelled. Potential enhancers (highlighted in yellow) were selected for knockout experiments to verify their regulatory function on the expression of target genes. **e.** The RT-qPCR analysis of the indicated gene expression levels in the DUX-induced GFP^+^ cells, including control (Ctrl) cells and different clones with homogeneous knockout of the potential enhancer regions highlighted in Fig. 4d. eKO: enhancer knockout. Control cells: cells transfected Cas9 plasmid with no gRNA. Values are means ± SD, n = 3 biological replicates. Two-sided paired t tests were performed. **f.** Schematic illustration of the procedure of nuclear transfer with control (Ctrl) mESCs or enhancer knockout (eKO) clones in Fig. 4e. Late 2-cell (L2C) embryos were collected for RT-qPCR analysis. **g.** The RT-qPCR analysis of the indicated gene expression levels in mouse L2C embryos generated from nuclear transfer with control (Ctrl) mESCs or enhancer knockout (eKO) clones in Fig. 4e (indicated with same color). Values are means ± SD, n = 3 biological replicates. Two-sided paired t tests were performed.

Next, we excised three enhancer regions located near the ZGA genes, *Ercc4* (Excision repair cross-complementing rodent repair deficiency, complementation group 4), *Stox1* (Storkhead box 1) and *Obox6* (Oocyte specific homeobox 6), to assess their functional roles (Fig. 4d). These three genes were not only highly expressed in DUX-induced GFP^+^ cells (Supplementary Fig. 7a), but also showed elevated expression during ZGA process (Supplementary Fig. 7b). For each enhancer region, a pair of sgRNAs was employed for the genomic deletion. Homozygous knockout (KO) clones derived from single cells for each enhancer were expanded and utilized for the further analysis (Supplementary Fig. 7c). RT-qPCR analysis revealed that deletion of these enhancers resulted in a significant reduction in the expression levels of their presumed target genes, *Ercc4*, *Stox1*, and *Obox6* in 2CLCs (Fig. 4e and Supplementary Fig. 7c). It is noted that the expression levels of *Dux* and *MERVL-Pol* remained unaffected, eliminating the possibility that the reduction in the expression of *Ercc4*, *Stox1*, and *Obox6* was due to a failure of Dux-mediated 2CLC induction (Fig. 4e).

To further investigate whether the deletion of the three enhancers could affect the expression of their target genes *in vivo*, one of the homozygous knockout clones of each enhancer was selected to conduct nuclear transfer (NT). NT-L2C embryos were subsequently collected for gene expression analysis (Fig. 4f). While the developmental rate of *Ercc4-*/*Stox1-*/*Obox6*-eKO 2C embryos had no apparent difference, the expression levels of *Ercc4* and *Stox1* were significantly reduced in their respective enhancer knockout (eKO) embryos, compared to the NT embryos derived from control (ctrl) cells (Fig. 4g and Supplementary Fig. 7d, e). Whereas, the expression of *Obox6* showed minimal changes (Fig. 4g). Considering that 2C embryos exhibit a notably higher expression level of *Obox6* compared to 2CLCs ^46^, we speculated that compensatory mechanisms, probably involving other unidentified enhancers, are responsible for regulating *Obox6* expression *in vivo* ^46^. These findings indicated that the enhancers identified in 2CLCs play an important role in regulating their target ZGA genes both *in vitro* and *in vivo*.

## Discussion

In this study, we conducted a comprehensive analysis of the entire mouse genome to pinpoint the functional enhancers in DUX-induced 2CLCs. Although 2CLCs partially mimic 2C embryos, our integrated analysis of epigenetic modification and transcriptome during ZGA has enabled us to shed light on the *cis*-regulatory elements governing gene expression during the ZGA process.

After fertilization, the quiescent zygotic genome entered into an open status without detectable topologically associated domains (TADs) ^2,33,34^. Previous studies suggested that the first zygotic transcription is a promiscuous process^48^. Enhancers might be dispensable for transcription in zygotes^35^ and become essential when chromatin is proposed to progressively adopt a repressive state during ZGA. Notably, in our data, we identified a group of promoters of minor ZGA genes, including *Dux*, *Zfp352*, *Zscan4*, have enhancer activities. Those enhancer-promoters have high levels of open chromatin-related histone modifications both i*n vitro* and *in vivo* and their corresponding genes show higher expression compared to the other transcripts. Moreover, the introduction of STARR constructs and fluorescence reporter plasmids containing those enhancer-promoter regions into embryos demonstrated their role as enhancers in 2C embryos. Considering that minor ZGA is critical for the proper ZGA process, probably by producing TFs to recruit Pol II to major ZGA-related targets ^4,49^. Rather than merely a promiscuous process, our results support the presence of an alternative mechanism which is independent of long-distance enhancers, facilitates a burst in gene expression during ZGA. Interestingly, the ZGA specific enhancer-promoters exclusively enriched binding motifs of both canonical transcription activator (including NRF1, YY1 and NFY) and 2C-specific TFs. Previous work showed that NFYA, a sequence-specific DNA binding subunit of NFY complex which forming a histone-like structure binding to DNA, promotes establishment of DNase I-hypersensitive site (DHS) in 2C embryos and is essential for the ZGA ^39,50,51^, suggesting that those enhancer-promoters might play a role in establishing of the chromatin structure in early embryos.

Mouse endogenous retrovirus elements play an important role in ZGA ^6,8,44,52^. Among them, MERVL is specifically activated at the 2C stage concomitant with ZGA and its *cis*-acting functions are required for the correct ZGA process ^8,44^. In our work, we offered the direct evidence of the enhancer function of MERVL elements in totipotent cells, and further characterized those enhancer-like MERVL elements. We revealed that the long terminal repeats (LTRs) of MERVL were the major elements which exhibited enhancer functions, occurring either as solo copies or as part of full-length MERVL. Although LTRs have conserved sequences, only around half of MERVL-LTRs exhibited the enhancer potential in 2CLCs. In total, 1115 annotated genes were associated with those LTR-enhancers, with 56.4% of them (629/1,115) upregulating in 2C::EGFP positive cells. As for *in vivo*, those LTRs also exhibited high levels of chromatin accessibility, with 315 associated annotated genes activated during the ZGA process. It should be noted that the other LTRs which hardly exhibited enhancer potential in 2CLCs, including those with STARR activities but low chromatin accessibility (MT2_Mm.C2), or those full-length LTRs exhibiting undetectable STARR activities (MT2_Mm.C3.1), were also bound by ZGA-critical TFs, DUX and OBOX, and exhibited an open chromatin state in 2C embryos. Further investigation is required to determine whether these LTRs perform an enhancer function *in vivo*. Collectively, our findings provide valuable insights into the *cis*-regulatory mechanisms governing the ZGA process both *in vitro* and *in vivo*.

## Methods

### Animals

All animal experiments were approved by Animal Ethics Committee of Anhui Medical University. B6D2F1 (C57BL/6×DBA2) adult mice were used in this study. The mice were kept in under controlled conditions, with a constant temperature and a 12-hour light/12-hour dark cycle.

### ESC culture and construction of the DUX-induced 2CLC model

The ES-E14Tg2a cells were cultured on 0.1% gelatin-coated (07903, Stem cell) plates with standard LIF/serum medium containing 15% FBS (10099141, Gibco), 1,000 U/mL mouse LIF (ESG1107, Millipore), 0.055 mM β-mercaptoethanol (21985023, Gibco), 0.1 mM non-essential amino acids (11140050, Gibco), 2 mM GlutaMAX (35050061, Gibco), 1 mM sodium pyruvate (11360070, Gibco), and 100 µg/mL penicillin/streptomycin (15140122, Gibco). The ESC lines were cultured in a 37 □ humidified incubator with 5% CO_2_. Medium was changed daily, and cells were routinely passaged every other day. The MERVL-LTR-EGFP-F2A-puroR reporter and doxycycline-inducible DUX overexpression cassette were cloned into PB-mCherry plasmid (Blasticidin resistance) and PB-3xFlag (Hygromycin resistance) respectively, and then transfected into E14Tg2a cells by Lipofectamine 3000 (L3000015, Invitrogen). ESCs with the 2C::EGFP reporter and doxycycline-inducible DUX overexpression cassette were selected simultaneously with blasticidin and hygromycin for one week. Single clones were then picked and expanded. Clone 3 which exhibited a high and stable proportion MERVL-EGFP positive cells upon doxycycline addition were chosen for the following experiments. 2 µg/mL doxycycline was used for the DUX induction.

### Karyotype analysis

The chromosome preparations were made in a previous protocol ^53^. When cells were grown with 60% confluence, 0.2 mg/mL colchicine (A600322, Sangon Biotech) was added into the medium and incubated at 37□ for 130 min (minutes). Then the cells were digested by 0.25% trypsin, and treated with 0.075 mol/L KCL (A501159, Sangon Biotech) at 37□ for 28 min. After centrifuge at 1000 rpm for 5 min, cells pellets were collected and incubated in fixative solution (methanol: glacial acetic acid=3:1) at 37□ for 40 min. The cell suspension was then dropped on the slide and oven baken at 75□ for 3 h. Subsequently, slides were incubated in the 0.25% trypsin for 10 s (seconds), and the trypsinization was terminated by saline solution. Slides were then stained by Giemsa dye solution for 10 min, and dried at room temperature (RT). Finally, Images were captured by Carl Zeiss imager Z2 (Carl Zeiss) and processed using Ikaros karyotyping system (MetaSystems).

### Flow cytometry

Cells were dissociated with 0.05% trypsin in 37□ for 5 min, then neutralized with the medium. After centrifuge at 1000 rpm for 5min, the cells were collected at the bottom of the tube and the supernatant was removed. The cells were resuspended in 300-400 µL PBS solution with 2% knockout serum (KSR, A3181501, Invitrogen) and 100 µg/mL penicillin/streptomycin (15140122, Gibco), then filtered with 35 mm strainer cap. Flow cytometry analysis was performed using the BD LSR Fortessa X-20 (BD), and cell sorting was performed on the BD FACSAria Fusion cell sorter (BD). Data and images were analysed and generated using FlowJo (V10) software.

### RNA isolation, RT-qPCR and total RNA-seq

Cellular total RNA was collected using a Trizol extraction method. The cells were lysed with 200 µL Trizol Reagent (15596026, Invitrogen) at RT for 5 min. The samples were added 40 µL chloroform and mixed well by vigorously upside down for 15 s, then put at RT for 15 min. After centrifuge at 14000 rpm, 4□ for 15 min, the upper aqueous solution was drawn into the new tube and add 100 µL cold isopropyl alcohol (I811925, Macklin), then mixed gently and placed on ice for 5-10 min. After centrifuge at 14000 rpm, 4□ for 10 min, the RNA was collected at the bottom of the tube and the supernatant was removed. Add 750 µL 75% ethanol (prepared with RNAase free water), and then reverse several times. After centrifuge at 14000 rpm, 4□ for 5 min, the RNA was collected at the bottom of the tube and the supernatant was removed. Dry the RNA at RT until the precipitate changed from white to transparent. Subsequently, the RNA was dissolved in 10-50 µL RNAase free water at 60□ for 10 min, and then the concentration of the RNA solution was measured. Complementary DNA was generated using PrimeScript II Reverse Transcriptase kit (2690A, TAKARA) and RT-qPCR was performed using TB Green PremixExTaq Kits (RR420, TAKARA) on QuantStudio^TM^ 7 Flex (ThermoFisher Scientific). Relative quantification was performed using the comparative CT method with normalization to GAPDH. Primers used for RT-qPCR are listed in Table S7.

The total RNA-seq library was prepared with rRNA depletion following stranded method. Briefly, the ribosomal RNA was depleted from total RNA using the Illumina Ribo-Zero plus rRNA Depletion kit (20037135, Illumina) following manufacturer’s instruction. RNA was then fragmented into 300∼350 bp fragments, and first strand cDNA was reverse-transcribed using fragmented RNA and dNTPs (dATP, dTTP, dCTP and dGTP). RNA was degraded using RNase H, and second strand cDNA was synthesised using DNA polymerase I and dNTPs (dATP, dUTP, dCTP and dGTP). Remaining overhangs of double-strand cDNA were converted into blunt ends via exonuclease/polymerase activities. After adenylation of 3’ ends of DNA fragments, sequencing adaptors were ligated to the cDNA and the library fragments were purified. The template without U was enriched by PCR, and the PCR product was purified to obtain the final library. Finally, the total RNA-seq library use PE150 (paired-end 150nt) sequencing (Illumina) at Berry Genomics Co., Ltd.

### 10× Genomics single-cell transcriptome sequencing (ScRNA-seq)

2C::EGFP reporter mESCs with (+Dox) or without (-Dox) *Dux* induction were digested and re-suspended in 0.5% BSA-PBS. Then 5,000-10,000 cells were collected under the microscope. The cells were counted using Countess (CellDropFL, Denvoix) and the concentration of single cell suspension was adjusted to 1,000 cells/µL. Cells were loaded according to standard protocol of the Chromium Next GEM Chip G Single Cell Kit (1000121, 10x Genomics) in order to capture 5,000 cells to 10,000 cells/chip position (V3 chemistry). All the remaining procedures including the library construction were performed according to the standard manufacturer’s protocol. Single cell libraries were sequenced on Illumina HiSeqXTen instruments using 150 nt paired-end sequencing at Berry Genomics Co., Ltd.

### Genome-Wide STARR-seq Plasmid pool Construction

The mouse genome was extracted from E14Tg2a mESCs and sonicated. Then sheared genomic DNA fragments of 700∼1200 bp were size selected on 1% agarose gel to construct the STARR-seq plasmid pool following the STARR-seq library cloning protocol ^47^. Illumina adapters were added to 5 μg of size-selected DNA fragments using NEBNext® Ultra™ II DNA Library Prep Kit for Illumina (NEB, #E7645S). The adapter-ligated fragments were amplified through 5 cycles across 10 reactions with library cloning primers ^47^, which added a 15nt sequence to the adapters for Gibson assembly. The PCR products were purified with 0.8× AMPure XP beads (A63881, Beckman) and then assembled into the Age1/Sal1-digested hSTARR-seq_ORI vector (99296, Addgene) following the Gibson Assembly Kit instructions (RK21020, ABclonal). The ligation products were purified with MinElute kit (28004, Qiagen) and transformed into MegaX DH10B™ T1R Electrocomp™ Cells (Invitrogen, #C640003). The Plasmid Plus Giga Kit (Qiagen, #12991) was used to extract the plasmid pool for screening.

### Genome-wide STARR-seq library preparation and sequencing

2C::EGFP reporter mESCs were pre-treated with 2 µg/mL doxycycline for 6 h, followed by 2 µg/mL doxycycline and 1 µg/mL puromycin for 18 h (as shown in Fig. 2a). Then 400 µg STARR plasmid pool was transfected into 400 million Dox-treated cells with Lipo3000 (L3000015, Invitrogen). After 16 hours of transfection, approximately 150 million cells were harvested. The STARR-seq libraries and input were generated according to a previous UMI-STARR-seq protocol ^47^. STARR-seq libraries were sequenced on Illumina HiSeqXTen instruments using 150 nt paired-end sequencing at Annoroad Co., Ltd.

### ULI-NChIP-seq

For ULI-NChIP-seq, 5,000 MERVL-GFP positive or MERVL-GFP negative cells were collected per reaction, and two replicates were performed for each stage. The cells were isolated and washed thoroughly in 0.5% BSA-PBS to avoid contamination. The ULI-NChIP procedure was performed as previously described ^54^. One microgram of histone H3K27ac (39133, Active Motif), H3K4me1 (5326S, Cell Signaling Technology) or H3K4me3 (9751S, Cell Signaling Technology) antibody was used for each immunoprecipitation reaction. The sequencing libraries were prepared using the NEBNext^®^ Ultra™ II DNA Library Prep Kit for Illumina^®^ (E7645S, NEB) and the NEBNext^®^ Multiplex Oligos for Illumina^®^ (Index Primers Set 1) (E7335S, NEB) following the manufacturer’s instructions. Paired-end sequencing with a 150 nt read length was performed on a NovaSeq (Illumina) platform at Berry Genomics Co., Ltd.

### ATAC-seq

For ATAC-seq, 50,000 MERVL-GFP positive or MERVL-GFP negative cells were collected per reaction, and two replicates were performed for each stage. The cells were isolated and washed thoroughly in 0.5% BSA-PBS to avoid contamination. The ATAC-seq libraries were generated according to a previous protocol with minor modifications ^2^. Briefly, samples were lysed in lysis buffer (10□mM Tris-HCl (pH 7.4), 10□mM NaCl, 3□mM MgCl_2_ and 0.1% NP-40) for 10□min on ice to prepare the nuclei. Immediately after lysis, nuclei were spun at 500*g* for 5□min to remove the supernatant. Then the nuclei were washed once with 200 µL 0.5% BSA-PBS and spun at 500*g* for 5□min to remove the supernatant. Nuclei were then incubated with the Tn5 transposome and tagmentation buffer at 37□°C for 30□min (TD502, Vazyme). After the tagmentation, the stop buffer was added directly into the reaction to end the tagmentation at RT for 5 min. The DNA was extracted using MinElute PCR Purification Kit (28004, QIAGEN) following the manufacturer’s instructions. PCR was performed to amplify the library for 13 cycles using the following PCR conditions: 72□°C for 5□min; 98□°C for 45□s; and thermocycling at 98□°C for 15□s, 60□°C for 30□s and 72□°C for 1□min; following by 72□°C for 5□min. After the PCR reaction, libraries were screened and purified with 0.65× and 1.8× AMPure XP beads (A63881, Beckman). Paired-end sequencing with a 150 nt read length was performed on a NovaSeq (Illumina) platform at Annoroad Co., Ltd.

### ULI-STARR-seq

The candidate enhancer and negative control sequences with Illumina adapters were synthesized and cloned into hSTARR-seq_ORI vector to construct the plasmid library (99296, Addgene). Then the STARR-seq plasmid library was purified and injected into the male pronucleus of zygotes (PN3-PN4, 20-21 h post-hCG injection). After 26h, the late 2-cell embryos (46-48h post-hCG injection) were lysed in hypotonic lysis buffer (S1550S, NEB) and the polyadenylated mRNAs were captured using the Magnetic mRNA Isolation Kit (S1550S, NEB). Then the sample was added URBO DNase (0.25 U /5 µL reaction system, AM2238, invitrogen) to digest genomic DNA at□37°C for 30□min. Subsequently, the polyadenylated mRNAs were purified using 1.8× RNA Clean beads (N412-02, Vazyme). The reverse transcription and subsequent steps followed the previous UMI-STARR-seq protocol ^47^. STARR-seq libraries were sequenced on Illumina HiSeqXTen instruments using 150 nt paired-end sequencing at Repugene Co., Ltd.

### Mouse zygote collection and Injection

Eight-to nine-week-old B6D2F1 females were superovulated by injecting 10 international units (IU) pregnant mare’s serum gonadotropin (PMSG), followed by 14 IU human chorionic gonadotropin (hCG) 48 h (hour) later. The superovulated B6D2F1 female mice mated with B6D2F1 adult males after hCG injection. Zygotes were collected from oviducts and cultured in KSOM medium (Millipore) at 37□ under 5% CO_2_. The successfully constructed plasmids were diluted to 20 ng/μl and microinjected into the pronuclei of mouse zygotes by using an Eppendorf FemtoJet 4i. The late-2-cell stage embryos were harvested 46-48 h after hCG injection.

Candidate enhancer and negative control plasmids were introduced by pronuclear injection, following the same procedure as described above. Purified candidate enhancer or negative control plasmids were co-injected with the mCherry reporter plasmid, with final concentrations of 4.5 ng/μl for the candidate enhancer or negative control constructs and 1.5 ng/μl for the mCherry plasmid. After pronuclear injection, zygotes were cultured under the same conditions, and fluorescence intensity was examined 24 hours later using a Nikon fluorescence microscope (Ti2-U).

### Nuclear transfer

All MII oocytes were collected from superovulated female B6D2F1 mice (8-9 weeks old) 14h post-hCG injection and removed the cumulus cells *in vitro* by 1% hyaluronidase. The collected MII oocytes were firstly transferred to droplets of HEPES-CZB medium containing 5 μg/ml cytochalasin B (CB). A piezo-assisted pipette with an inner diameter of 8 μm was then used to pierce zona pellucida and aspirate the chromatin spindle region with a minimum volume of oocyte cytoplasm. The enucleated oocytes were thoroughly washed and preincubated at 37□ in KSOM medium with amino acids until nuclear transfer.

The mESCs were pretreated by culturing in medium containing 0.05 μg/ml demecolcine (Sigma) for 12 h to arrest the mESCs at M phase. Then pretreated mESCs were digested with 0.05% trypsin-EDTA, resuspended the cells in HEPES-CZB and stored at 4□ before injection. The pretreated mESCs were gently injected into the enucleated oocytes. Around 1 h after nuclear transfer, the reconstructed embryos were activated for 6 h by Ca^2+^-free CZB containing 5 nM trichostatin A (TSA), 5 μg/ml CB and 10 mM SrCl_2_. Following activation, the reconstructed embryos were transferred into KSOM medium with amino acids containing 5nM TSA for 4 h. Then all of the reconstructed embryos were subsequently cultured in KSOM medium with amino acids at 37 □ under 5% CO_2_. Late 2-cell embryos were collected at 31.5-32 h (including 6 h activation) for the RT-qPCR.

### Enhancer knockout (eKO) in mESCs

The eKO mESCs were performed using CRISPR-Cas9 genome editing with a pair of guide RNAs (gRNAs) for each enhancer region. Online tool (Benchling) was used to design gRNAs. The top listed gRNAs were then cloned into the pSpCas9(BB)-2A-Puro (PX459) V2.0 vector (62988, Addgene) (Table S7, sgRNAs). Similar to the above description, MERVL-LTR-EGFP reporter and Dox-inducible Dux expression cassette were transfected into E14Tg2a cells to generate the Dox-inducible Dux overexpression 2C::EGFP reporter mESCs without puromycin resistance (inducible Dux mCherry-MERVL-EGFP clone 28, iDux-mcmg-Clone 28). The 2C::EGFP reporter mESCs without puromycin resistance were then transfected with the Cas9/gRNA plasmids. Subsequently, Single cells were sorted after 2-day treatment with 1.5μg/ml puromycin and amplified. Then the genomic DNA from single cell-derived clones was extracted for genotyping. Primers used for genotyping are listed in Table S7. Homogenous eKO clones were used for the following analysis.

### Bioinformatics analysis

#### Single-cell RNA-seq analysis

Single-cell RNA-seq reads were mapped to the mouse reference genome (mm10) using STAR (v2.7.6a). Cells were filtered based on a minimum threshold of 1500 expressed genes per cell, resulting in a total of 4961 retained cells: 3209 cells with Dox (+Dox) and 1752 cells without Dox (-Dox). Subsequently, single-cell data integration was performed using Seurat’s canonical correlation analysis (CCA) integration tool (Seurat v4.0.4) with the FindIntegrationAnchors and IntegrateData functions. The data from this study and the scRNA-seq data of the in-house mouse preimplantation embryos, published data sets from GSE121459 (mESCs, intermediate cells, and 2C-like cells), GSE168728 (totipotent blastomere-like cells (TBLCs) and pluripotent stem cells (PSCs)), and GSE185005 (chemically induced totipotent stem cells, ciTotiSCs) were scaled using 2000 highly variable genes, and principal component analysis (PCA) was conducted using 30 principal components to compute UMAP coordinates with RunUMAP. Cell clusters were identified through a shared nearest neighbor (SNN) modularity optimization algorithm using the FindClusters function.

#### Total RNA-seq data analysis

The raw RNA-seq reads were processed using TrimGalore (v.0.6.4) (https://github.com/ FelixKrueger/TrimGalore) to remove adapter sequences and low-quality bases. The cleaned reads were then aligned to the mouse Gencode (vM15) transcriptome with RSEM (version 1.2.22) ^55^. Gene expression values were normalized across samples using the variance modeling tool DESeq2 (version 1.30.0) ^56^. Differentially expressed genes were obtained using DESeq2 (version 1.30.0) ^56^. Gene ontology analysis was performed using clusterProfiler (version 3.18.0) ^57^.

#### ChIP-seq and ATAC-seq data analysis

Analysis of ATAC-seq and ChIP-seq data were performed essentially as previously described^58^. The raw reads were processed using TrimGalore (v.0.6.4) to remove adapter sequences and low-quality bases. The cleaned reads were aligned to the mouse mm10 genome using bowtie2 (v2.2.5) ^59^, with the options: ‘‘-p 10 –very-sensitive –end-to-end –no-unal –phred33 –no-mixed -X 2000’’. To analyze repetitive sequences, the multi-mapped reads were kept but only the best alignment was reported; if more than one equivalent best alignment was found, then one random alignment was reported. The PCR duplicates were removed by Picard (https://broadinstitute.github.io/picard/) with the MarkDuplicates function. Peaks were called by MACS3 ^60^ by default settings. Alignment bam files were transformed into read coverage files (bigwig format) using deepTools (v2.5.4) ^61^ with the RPKM (Reads Per Kilobase per Million mapped reads) normalization method. Coordinates and annotations of TEs were downloaded from the UCSC Genome Browser (GRCm38/mm10) version of RepeatMasker (http://hgdownload.soe.ucsc.edu/goldenPath/mm10/database/rmsk.txt.gz). Heatmaps and pileups were generated using deepTools. Motif discoverywere performed by HOMER^62^ with the command:”findMotifsGenome.pl input.bed mm10 output -p 20 -size given”. Peaks annotation was performed by HOMER with the command “annotatePeaks.pl input.bed mm10”.

#### STARR-seq data analysis

The raw reads of STARR-seq and input were processed using TrimGalore (v.0.6.4) to remove adapter sequences and low-quality bases. The cleaned reads were aligned to the mouse mm10 genome using bowtie2 (v2.2.5) ^59^, with the options: -p 48 --very-sensitive --end-to-end --no-unal --phred33 -X 2000’. To analyze repetitive sequences, the multi-mapped reads were kept but only the best alignment was reported; if more than one equivalent best alignment was found, then one random alignment was reported. The PCR duplicates were removed by Picard (https://broadinstitute.github.io/picard/) with the MarkDuplicates function. Alignment bam files were transformed into read coverage files (bigwig format) using deepTools (v2.5.4) ^61^ with the RPKM normalization method. Genome coverage was computed with deeptools using the pybigwig function. Enriched genomic peaks were called using a 500bp sliding window algorithm, with a sliding step of 250 bp, using custom scripts. For every 500bp window in the genome, the number of reads falling in that window for a given sample (normalized by RPKM method) was divided by the number of reads in that window for the corresponding input file (also normalized by RPKM method). Enriched regions were initially formed by taking all 500 bp windows whose signal-over-input value exceeds a 2-fold threshold. Finally, any remaining overlaps among regions were merged, to produce the final peak calls. To identify the differential STARR-seq signal between 2CLCs and ESCs, the peaks from those two samples were merged using the mergeBed command from bedtools ^62^. Then deeptools’ pybigwig was used to calculate the value for each region. Genome coverage was computed with deeptools using the pybigwig function. Regions where the value differed by more than 2-fold between 2CLCs and ESCs were considered to be differential peaks. Motif discovery were performed by HOMER^62^ with the command:”findMotifsGenome.pl input.bed mm10 output -p 20 -size given”. Peaks annotation were performed by HOMER with the command “annotatePeaks.pl input.bed mm10”. Heatmap were made by deeptools^61^. The CpG island (CGI) annotation were downloaded from UCSC browser.

#### ULI-STARR-seq data analysis

First, we constructed a genome containing only our target sequences of interest. We then mapped the cleaned sequencing data back to this reference genome using Bowtie2 with the options: -p 48 --very-sensitive --end-to-end --no-unal --phred33 -X 2000’. Reads that aligned to identical genomic coordinates were identified as PCR duplicates, and only one representative read from each set of duplicates was retained for subsequent analysis. Samtools was used to count the number of sequencing reads mapped to each reference sequence. The ratio of reads mapped to a reference sequence in embryos compared to the input was considered the STARR-seq signal value for that sequence in the embryos. A reference sequence was considered enriched for STARR-seq signal in 2C embryos if the number of reads mapped to that sequence exceeded 1.5 times the average of the negative control.

## Data and code availability

High-throughput sequencing data generated in this study are available in the Gene Expression Omnibus with accession number GSE283389 (token: qfubikeollujjuz). The scRNA-seq data were from GSE121459 ^16^, GSE168728 ^20^ and GSE185005 ^19^. The ATAC-seq and RNA-seq data for pre-implantation embryos were from GSE66390 ^2^. The H3K4me1, H3K4me3 and H3K27ac ChIP-seq data for pre-implantation embryos were from GSE179941, GSE217970 ^23^, GSE71434 ^22^. The Obox ChIP-seq data were from GSE215813 ^46^, the Dux ChIP-seq data were from GSE95517 ^9^. This paper does not report original code.

## Acknowledgments

We would like to thank Zhe Xu and Haiyu Wang from the Man Zhang laboratory for help with the experiments and bioinformatics analysis. We also thank the Animal Ethics Committee of Anhui Medical University for their support. Furthermore, we are grateful to Dr. Xiaoying Fan for her critical reading and our laboratory colleagues for their assistance with experiments and advice. This work was supported by grants to M.Z. from the National Natural Science Foundation of China (32570958), the Pearl River Talent Recruitment Program (2021ZT09Y233), and Major Project of Guangzhou National Laboratory (grant no. GZNL2023A02005 and GZNL2024A02005); grants to J.H. from the National Key R&D Program of China (2021YFA1102200), the National Natural Science Foundation of China (32570664, 32370856), the Guangdong Natural Science Funds for Distinguished Young Scholar (2023B1515020111) and grant to T.P. from National Natural Science Foundation of China (grant no. 32400600).

## Author contributions

M.Z. conceived the project and designed the experiments. L.Y., T.P. performed most of the experiments, with the help of B.H. and L.L., D.L., T.Z. and Y.Z., performed embryo injection experiments. J.H. performed most of the bioinformatic data analysis, with the help of X.Z., L.Y., T.P. and M.Z., L.Y., T.P., J.H. and J.C. wrote the manuscript.

## Supplementary information

Supplementary information, including seven tables, can be found with this article.

**Supplementary Table 1. The differentially expressed genes (DEGs) between MERVL-GFP positive and MERVL-GFP negative cells. Related to Fig. 1**. (see attached the excel file named “Table S1. The DEGs between GFP+ and GFP-cells”)

**Supplementary Table 2. List of defined ZGA genes. Related to Fig. 2**. (see attached the excel file named “Table S2. List of defined ZGA genes”)

**Supplementary Table 3. List of the ZGA genes with enhancer activities at promoter regions. Related to Fig. 2**. (see attached the excel file named “Table S3. List of the ZGA genes with enhancer-promoters”)

**Supplementary Table 4. Transposable elements (TEs) enriched at 2C-specific (2C.spe) or ESC-specific (ESC.spe) STARR peaks. Related to Fig. 3**. (see attached the excel file named “Table S4. TEs enriched at 2C.spe or ESC.spe STARR peaks”)

**Supplementary Table 5. The relevant analysis of MERVL.C1 associated genes. Related to Fig. 3**. (see attached the excel file named “Table S5. Relevant analysis of MERVL.C1 associated genes”)

**Supplementary Table 6. Relevant information of synthesized enhancer candidates and negative controls for ULI-STARR-seq. Related to Fig. 4**. (see attached the excel file named “Table S6. Synthesized sequences for ULI-STARR-seq”)

**Supplementary Table 7. The list of oligos, including sgRNA, primers for Genotyping and primers for RT-qPCR. Related to methods.** (see attached the excel file named “Table S7. The list of oligos”)

## References

1 McGrath, J. & Solter, D. Completion of mouse embryogenesis requires both the maternal and paternal genomes. Cell 37, 179–183 (1984). 10.1016/0092-8674(84)90313-1

2 Wu, J. et al. The landscape of accessible chromatin in mammalian preimplantation embryos. Nature 534, 652–657 (2016). 10.1038/nature18606

3 Eckersley-Maslin, M. A., Alda-Catalinas, C. & Reik, W. Dynamics of the epigenetic landscape during the maternal-to-zygotic transition. Nature Reviews Molecular Cell Biology 19, 436–450 (2018). 10.1038/s41580-018-0008-z

4 Abe, K.-i., et al. Minor zygotic gene activation is essential for mouse preimplantation development. Proceedings of the National Academy of Sciences 115 (2018). 10.1073/pnas.1804309115

5 Jukam, D., Shariati, S. A. M. & Skotheim, J. M. Zygotic Genome Activation in Vertebrates. Dev Cell 42, 316–332 (2017). 10.1016/j.devcel.2017.07.026

6 Macfarlan, T. S. et al. Embryonic stem cell potency fluctuates with endogenous retrovirus activity. Nature 487, 57–63 (2012). 10.1038/nature11244

7 Peaston, A. E. et al. Retrotransposons regulate host genes in mouse oocytes and preimplantation embryos. Dev Cell 7, 597–606 (2004). 10.1016/j.devcel.2004.09.004

8 Sakashita, A. et al. Transcription of MERVL retrotransposons is required for preimplantation embryo development. Nat Genet 55, 484–495 (2023). 10.1038/s41588-023-01324-y

9 Hendrickson, P. G. et al. Conserved roles of mouse DUX and human DUX4 in activating cleavage-stage genes and MERVL/HERVL retrotransposons. Nat Genet 49, 925–934 (2017). 10.1038/ng.3844

10 Ishiuchi, T. et al. Early embryonic-like cells are induced by downregulating replication-dependent chromatin assembly. Nat Struct Mol Biol 22, 662–671 (2015). 10.1038/nsmb.3066

11 Eckersley-Maslin, Mélanie A. et al. MERVL/Zscan4 Network Activation Results in Transient Genome-wide DNA Demethylation of mESCs. Cell Reports 17, 179–192 (2016). 10.1016/j.celrep.2016.08.087

12 Iturbide, A. et al. Retinoic acid signaling is critical during the totipotency window in early mammalian development. Nat Struct Mol Biol (2021). 10.1038/s41594-021-00590-w

13 De Iaco, A. et al. DUX-family transcription factors regulate zygotic genome activation in placental mammals. Nat Genet 49, 941–945 (2017). 10.1038/ng.3858

14 Eckersley-Maslin, M. et al. Dppa2 and Dppa4 directly regulate the Dux-driven zygotic transcriptional program. Genes & Development 33, 194–208 (2019). 10.1101/gad.321174.118

15 Hu, Z. et al. Maternal factor NELFA drives a 2C-like state in mouse embryonic stem cells. Nat Cell Biol 22, 175–186 (2020). 10.1038/s41556-019-0453-8

16 Fu, X., Wu, X., Djekidel, M. N. & Zhang, Y. Myc and Dnmt1 impede the pluripotent to totipotent state transition in embryonic stem cells. Nat Cell Biol 21, 835–844 (2019). 10.1038/s41556-019-0343-0

17 Xu, Y. et al. Derivation of totipotent-like stem cells with blastocyst-like structure forming potential. Cell Res 32, 513–529 (2022). 10.1038/s41422-022-00668-0

18 Yang, M. et al. Chemical-induced chromatin remodeling reprograms mouse ESCs to totipotent-like stem cells. . Cell Stem Cell 29, 1–19 (2022). 10.1016/j.stem.2022.01.010

19 Hu, Y. et al. Induction of mouse totipotent stem cells by a defined chemical cocktail. Nature 617, 792–797 (2023). 10.1038/s41586-022-04967-9

20 Shen, H. et al. Mouse totipotent stem cells captured and maintained through spliceosomal repression. Cell 184, 2843–2859 e2820 (2021). 10.1016/j.cell.2021.04.020

21 Long, H. K., Prescott, S. L. & Wysocka, J. Ever-Changing Landscapes: Transcriptional Enhancers in Development and Evolution. Cell 167, 1170–1187 (2016). 10.1016/j.cell.2016.09.018

22 Zhang, B. et al. Allelic reprogramming of the histone modification H3K4me3 in early mammalian development. Nature 537, 553–557 (2016). 10.1038/nature19361

23 Liu, B. et al. Mapping putative enhancers in mouse oocytes and early embryos reveals TCF3/12 as key folliculogenesis regulators. Nat Cell Biol 26, 962–974 (2024). 10.1038/s41556-024-01422-x

24 Muerdter, F., Boryn, L. M. & Arnold, C. D. STARR-seq -principles and applications. Genomics 106, 145–150 (2015). 10.1016/j.ygeno.2015.06.001

25 Barakat, T. S. et al. Functional Dissection of the Enhancer Repertoire in Human Embryonic Stem Cells. Cell Stem Cell 23, 276–288 e278 (2018). 10.1016/j.stem.2018.06.014

26 Huang, B. et al. Inhibition of HDAC activity directly reprograms murine embryonic stem cells to trophoblast stem cells. Developmental Cell (2024). 10.1016/j.devcel.2024.05.009

27 Muerdter, F. et al. Resolving systematic errors in widely used enhancer activity assays in human cells. Nat Methods 15, 141–149 (2018). 10.1038/nmeth.4534

28 Peng, T. et al. STARR-seq identifies active, chromatin-masked, and dormant enhancers in pluripotent mouse embryonic stem cells. Genome Biol 21, 243 (2020). 10.1186/s13059-020-02156-3

29 Andersson, R. & Sandelin, A. Determinants of enhancer and promoter activities of regulatory elements. Nat Rev Genet 21, 71–87 (2020). 10.1038/s41576-019-0173-8

30 Shlyueva, D., Stampfel, G. & Stark, A. Transcriptional enhancers: from properties to genome-wide predictions. Nature Reviews Genetics 15, 272–286 (2014). 10.1038/nrg3682

31 Whiddon, J. L., Langford, A. T., Wong, C. J., Zhong, J. W. & Tapscott, S. J. Conservation and innovation in the DUX4-family gene network. Nat Genet 49, 935–940 (2017). 10.1038/ng.3846

32 Choi, Y. J. et al. Deficiency of microRNA miR-34a expands cell fate potential in pluripotent stem cells. Science 355 (2017). 10.1126/science.aag1927

33 Du, Z. et al. Allelic reprogramming of 3D chromatin architecture during early mammalian development. Nature 547, 232–235 (2017). 10.1038/nature23263

34 Ke, Y. et al. 3D Chromatin Structures of Mature Gametes and Structural Reprogramming during Mammalian Embryogenesis. Cell 170, 367–381 e320 (2017). 10.1016/j.cell.2017.06.029

35 Majumder, S., Miranda, M. & DePamphilis, M. L. Analysis of gene expression in mouse preimplantation embryos demonstrates that the primary role of enhancers is to relieve repression of promoters. EMBO J 12, 1131–1140 (1993). doi: 10.1002/j.1460-2075.1993.tb05754.x.

36 Zabidi, M. A. et al. Enhancer-core-promoter specificity separates developmental and housekeeping gene regulation. Nature 518, 556–559 (2015). 10.1038/nature13994

37 Hounkpe, B. W., Chenou, F., de Lima, F. & De Paula, Erich V. HRT Atlas v1.0 database: redefining human and mouse housekeeping genes and candidate reference transcripts by mining massive RNA-seq datasets. Nucleic Acids Research 49, D947–D955 (2021). 10.1093/nar/gkaa609

38 Virbasius, C. A., Virbasius, J. V. & Scarpulla, R. C. NRF-1, an activator involved in nuclear-mitochondrial interactions, utilizes a new DNA-binding domain conserved in a family of developmental regulators. Genes Dev 7, 2431–2445 (1993). 10.1101/gad.7.12a.2431

39 Lu, F. et al. Establishing Chromatin Regulatory Landscape during Mouse Preimplantation Development. Cell 165, 1375–1388 (2016). 10.1016/j.cell.2016.05.050

40 Duttke, S. H. et al. Position-dependent function of human sequence-specific transcription factors. Nature 631, 891–898 (2024). 10.1038/s41586-024-07662-z

41 Grow, E. J. et al. p53 convergently activates Dux/DUX4 in embryonic stem cells and in facioscapulohumeral muscular dystrophy cell models. Nature Genetics 53, 1207–1220 (2021). 10.1038/s41588-021-00893-0

42 Zhou, C., Wang, M., Zhang, C. & Zhang, Y. The transcription factor GABPA is a master regulator of naive pluripotency. Nature Cell Biology 27, 48–58 (2025). 10.1038/s41556-024-01554-0

43 Yang, F. et al. DUX-miR-344-ZMYM2-Mediated Activation of MERVL LTRs Induces a Totipotent 2C-like State. Cell Stem Cell 26, 234–250 e237 (2020). 10.1016/j.stem.2020.01.004

44 Yang, J., Cook, L. & Chen, Z. Systematic evaluation of retroviral LTRs as cis-regulatory elements in mouse embryos. Cell Reports 43 (2024). 10.1016/j.celrep.2024.113775

45 Kigami, D., Minami, N., Takayama, H. & Imai, H. MuERV-L is one of the earliest transcribed genes in mouse one-cell embryos. Biol Reprod 68, 651–654 (2003). 10.1095/biolreprod.102.007906

46 Ji, S. et al. OBOX regulates mouse zygotic genome activation and early development. Nature 620, 1047–1053 (2023). 10.1038/s41586-023-06428-3

47 Neumayr, C., Pagani, M., Stark, A. & Arnold, C. D. STARR-seq and UMI-STARR-seq: Assessing Enhancer Activities for Genome-Wide-, High-, and Low-Complexity Candidate Libraries. Curr Protoc Mol Biol 128, e105 (2019). 10.1002/cpmb.105

48 Abe, K. et al. The first murine zygotic transcription is promiscuous and uncoupled from splicing and 3’ processing. EMBO J 34, 1523–1537 (2015). 10.15252/embj.201490648

49 Liu, B. et al. The landscape of RNA Pol II binding reveals a stepwise transition during ZGA. Nature 587, 139–144 (2020). 10.1038/s41586-020-2847-y

50 Oldfield, Andrew J. et al. Histone-Fold Domain Protein NF-Y Promotes Chromatin Accessibility for Cell Type-Specific Master Transcription Factors. Molecular Cell 55, 708–722 (2014). 10.1016/j.molcel.2014.07.005

51 Nardini, M. et al. Sequence-Specific Transcription Factor NF-Y Displays Histone-like DNA Binding and H2B-like Ubiquitination. Cell 152, 132–143 (2013). 10.1016/j.cell.2012.11.047

52 Li, X. et al. LINE-1 transcription activates long-range gene expression. Nature Genetics 56, 1494–1502 (2024). 10.1038/s41588-024-01789-5

53 Yunis, J. J., Roldan, L., Yasmineh, W. G. & Lee, J. C. Staining of satellite DNA in metaphase chromosomes. Nature 231, 532–533 (1971). 10.1038/231532a0

54 Liu, X. et al. Distinct features of H3K4me3 and H3K27me3 chromatin domains in pre-implantation embryos. Nature 537, 558–562 (2016). 10.1038/nature19362

55 Li, B. & Dewey, C. N. RSEM: accurate transcript quantification from RNA-Seq data with or without a reference genome. BMC Bioinformatics 12, 323 (2011). 10.1186/1471-2105-12-323

56 Love, M. I., Huber, W. & Anders, S. Moderated estimation of fold change and dispersion for RNA-seq data with DESeq2. Genome Biol 15, 550 (2014). 10.1186/s13059-014-0550-8

57 Yu, G., Wang, L. G., Han, Y. & He, Q. Y. clusterProfiler: an R package for comparing biological themes among gene clusters. Omics 16, 284–287 (2012). 10.1089/omi.2011.0118

58 Wu, F. et al. Species-specific rewiring of definitive endoderm developmental gene activation via endogenous retroviruses through TET1-mediated demethylation. Cell Rep 41, 111791 (2022). 10.1016/j.celrep.2022.111791

59 Langmead, B. & Salzberg, S. L. Fast gapped-read alignment with Bowtie 2. Nat Methods 9, 357–359 (2012). 10.1038/nmeth.1923

60 Zhang, Y., et al. Model-based analysis of ChIP-Seq (MACS). Genome Biol 9, R137 (2008). 10.1186/gb-2008-9-9-r137

61 Ramírez, F. et al. deepTools2: a next generation web server for deep-sequencing data analysis. Nucleic Acids Res 44, W160–165 (2016). 10.1093/nar/gkw257

62 Heinz, S. et al. Simple combinations of lineage-determining transcription factors prime cis-regulatory elements required for macrophage and B cell identities. Mol Cell 38, 576–589 (2010). 10.1016/j.molcel.2010.05.004

