## Supplementary information for "High-throughput functional characterization of enhancers in totipotent-like cells"

#### Supplementary Fig. 1

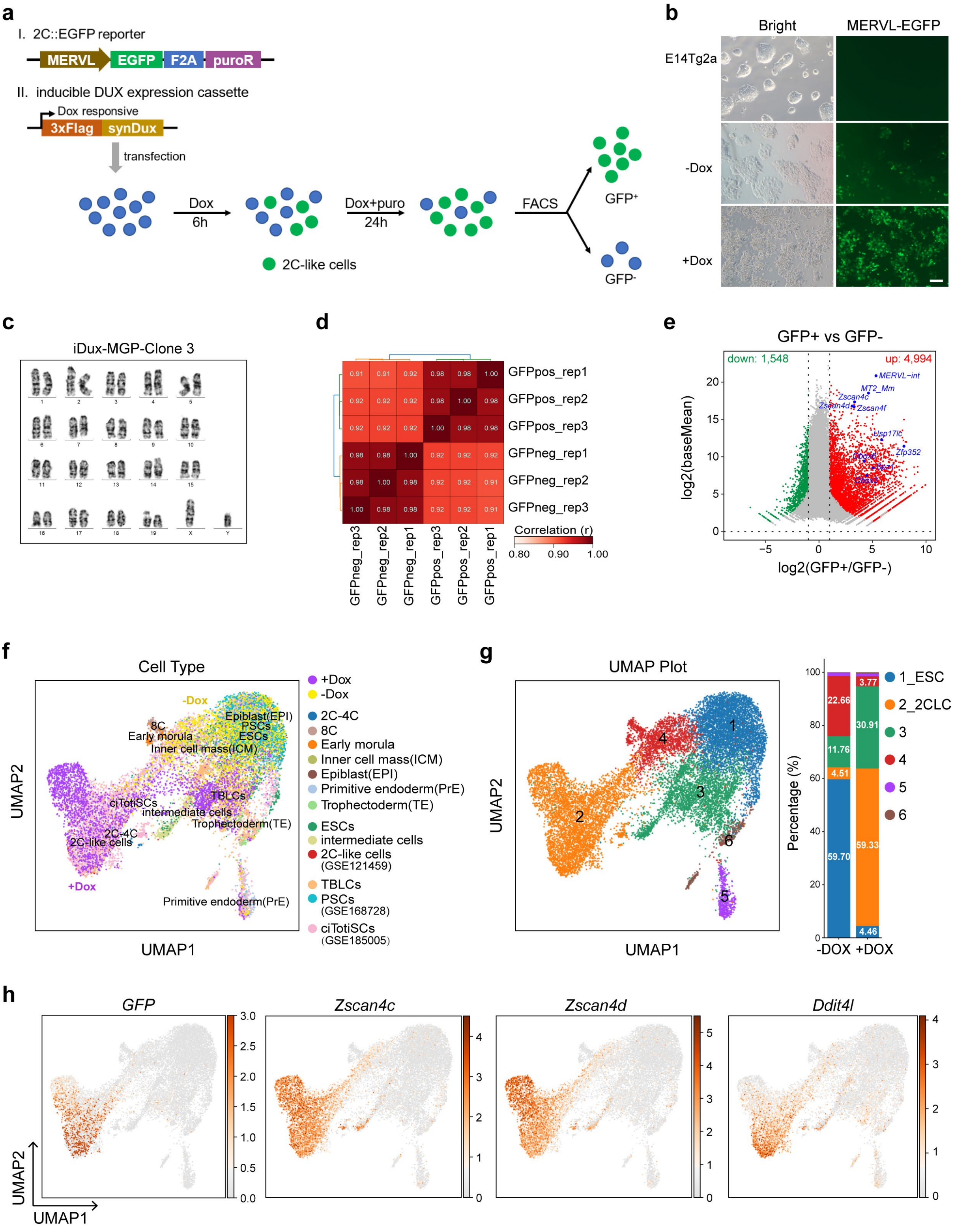

**Supplementary Fig. 1 | Establishment of a DUX-induced 2CLC model. Related to Fig. 1.**

**(a)** Schematic illustration of inducing 2C-like cells (2CLCs) from doxycycline (Dox)-inducible DUX overexpression MERVL-EGFP reporter cell lines. Enhanced green florescence protein (EGFP) and Puromycin resistant (puroR) coding sequences are under the control of the LTR of MERVL. syn*Dux* refers to codon-optimized exogenous *Dux*.

**(b)** Images showing wildtype E14Tg2a cells and inducible DUX overexpression MERVL-EGFP (2C::EGFP) cells with (+) or without (-) doxycycline (Dox) in brightfield and green fluorescence channel. Scale bar: 100 μm.

**(c)** Karyotype analysis of iDux-MGP-Clone 3 cell line. iDux-MGP-Clone 3: single cell derived doxycycline-inducible DUX overexpression, MERVL-EGFP-F2A-puroR clone 3 cell line.

**(d)** Heatmap showing the correlation between the total RNA-seq biological replicates of MERVL-$GFP^-$ and MERVL-$GFP^+$ cell samples. n = 3 biological replicates.

**(e)** M-versus-A (MA) plot showing the differential expressed genes and transposable elements (TEs) between MERVL-$GFP^+$ and MERVL-$GFP^-$ cells.

**(f)** Uniform manifold approximation and projection (UMAP) analysis of single cell RNA-seq data from inducible DUX overexpression 2C::EGFP reporter mESCs (as shown in Supplementary Fig. 1a) with (+) or without (-) 2 μg/mL doxycycline (Dox), in-house mouse preimplantation embryos, and published data sets from GSE121459 (mESCs, intermediate cells, and 2C-like cells), GSE168728 (totipotent blastomere-like cells (TBLCs) and pluripotent stem cells (PSCs)), and GSE185005 (chemically induced totipotent stem cells, ciTotiSCs). -Dox: Dox-inducible DUX overexpression 2C::EGFP reporter mESCs without doxycycline treatment. +Dox: Dox-inducible DUX overexpression 2C::EGFP reporter mESCs with doxycycline treatment.

**(g)** UMAP plot for scRNA-seq analysis of the cells from Supplementary Fig. 1f, showing the 6 clusters (left). The percentage of -Dox group and the +Dox group in the 6 main clusters are shown (right). -Dox: Dox-inducible DUX overexpression 2C::EGFP reporter mESCs without doxycycline treatment. +Dox: Dox-inducible DUX overexpression 2C::EGFP reporter mESCs with doxycycline treatment.

**(h)** UMAP plot showing the expression of *EGFP* and 2C markers (*Zscan4c*, *Zscan4d* and *Ddit4l*) in the 6 clusters in Supplementary Fig. 1g.

#### Supplementary Fig. 2

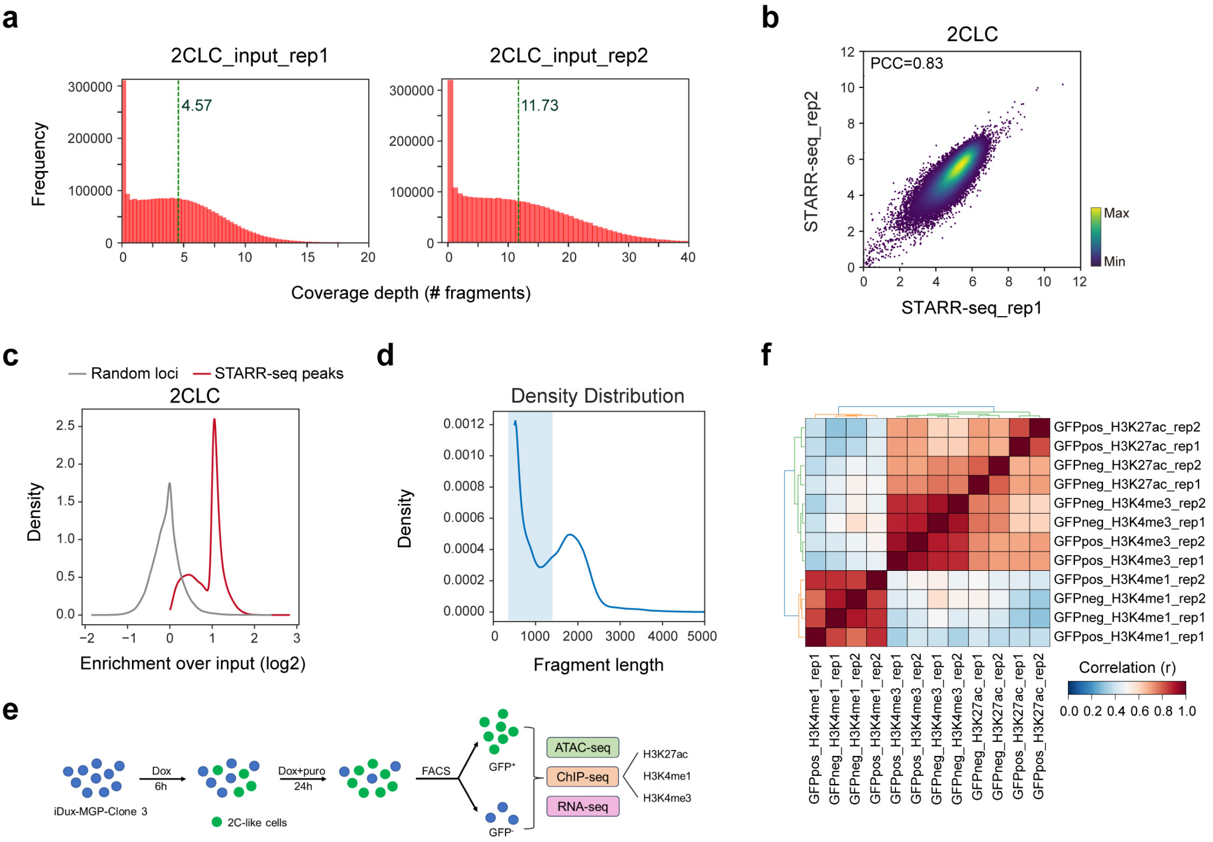


**Supplementary Fig. 2 | Quality assessment of high-throughput sequencing data. Related to Fig. 1.**

**(a)** Genome wide coverage and depth per STARR-seq input library.

**(b)** The correlation between the whole-genome SATRR-seq biological replicates of 2CLC samples. PCC: Pearson's correlation coefficient. n = 2 biological replicates.

**(c)** STARR-seq enrichment analysis at STARR-seq peaks compared to randomly selected length-matched regions. A 2-fold (log2(fold change) ≥ 1) or higher STARR-seq enrichment over input was observed in less than 0.7% (308/45,917) randomly selected regions and used as a cutoff for defining significant enhancer activity.

**(d)** Fragment length distribution of 2CLC STARR-seq peaks.

**(e)** Schematic illustration of the ATAC-seq, ChIP-seq and total RNA-seq sample collections.

**(f)** The correlation between the ChIP-seq biological replicates of MERVL-GFP$^-$ and MERVL-GFP$^+$ cells. n = 2 biological replicates.

### Supplementary Fig. 3

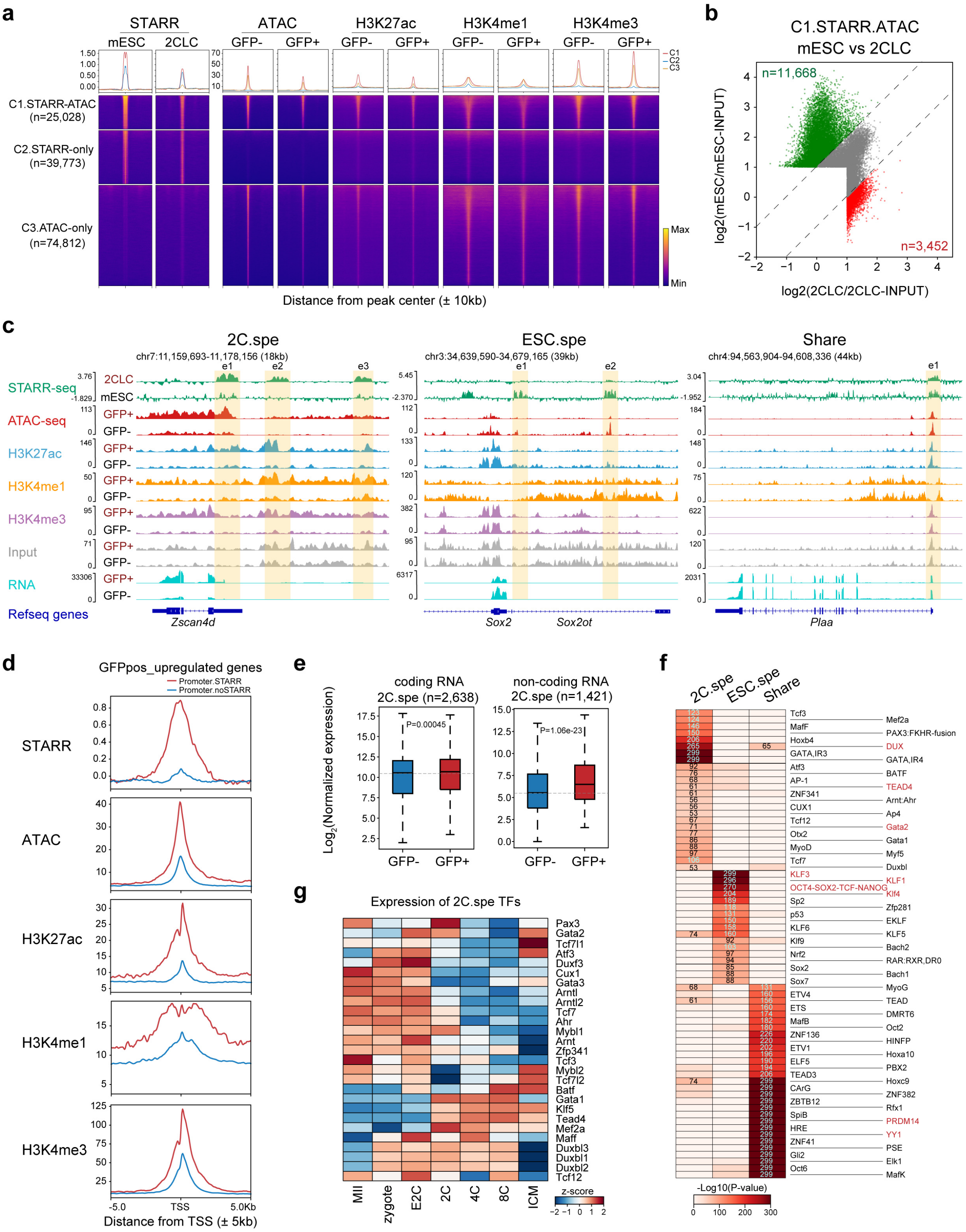

**Supplementary Fig. 3 | The characteristic analysis of enhancers identified by STARR-seq in 2CLCs and ESCs. Related to Fig. 1.**

**(a)** Heatmap of STARR-seq signals in mESCs and 2CLCs, ATAC-seq and ChIP-seq signals in MERVL-GFP$^-$ and MERVL-GFP$^+$ cells. Signals were computed on the STARR- (C1 and C2) or ATAC-seq peak (C3) flanked by 10 kb. Regions were clustered by class and ranked by decreasing STARR-seq signal (log2(RPKM)) in mESCs. C1.STARR-ATAC: the regions with both STARR-seq and ATAC-seq signals in mESCs or 2CLCs; C2.STARR-only: the regions with only STARR-seq signals but no ATAC-seq signals in mESCs or 2CLCs; C3.ATAC-only: the regions with no STARR-seq signals and only ATAC-seq signals in mESCs or 2CLCs. The peak number of each cluster is indicated.

**(b)** Scatter plot illustrating the STARR-seq enrichment in 2CLCs or ESCs within C1 STARR peaks (in Supplementary Fig. 3a). Differential peaks that are elevated in 2CLCs are marked in red (n = 3,452), while those elevated in mESCs are shown in green (n = 11,668). Non-differential peaks are represented in gray. The analysis was conducted using a fold change (FC) threshold of ≥ 2 with $p < 0.05$.

**(c)** Genome browser view showing selected 2C-specific (2C.spe, left), ESC-specific (ESC.spe, middle), and 2CLC/ESC shared (share, right) enhancer loci. The STARR-seq track depicts the enrichment over input. The highlighted regions were regarded as potential enhancers of indicated genes.

**(d)** Line chart showing STARR-seq, ATAC-seq, H3K27ac, H3K4me1 and H3K4me3 signals at the indicated promoter regions (TSS ± 5 kb) of upregulated genes in MERVL-GFP$^+$ cells, categorized by the presence (STARR) or absence (noSTARR) of STARR-seq signals in 2CLCs.

**(e)** Boxplot showing the expression levels of the transcripts flanked by 2C-specific (2C.spe) STARR peaks within 20 kb (± 20kb) in MERVL-GFP$^-$ and MERVL-GFP$^+$ cells. The transcripts were displayed as coding and long non-coding genes. P-values were from a Mann-Whitney U test.

**(f)** Heatmap showing transcription factor (TF) motifs identified from 2C-specific, ESC-specific and share STARR peaks from Fig. 2b. The color indicated the enrichment of the motif (-log10 P-value).

**(g)** Heatmap showing the RNA expression level of the TFs that its motifs were enriched in 2C-specific STARR peaks (from Supplementary Fig. 3f) during mouse embryonic development.

#### Supplementary Fig. 4

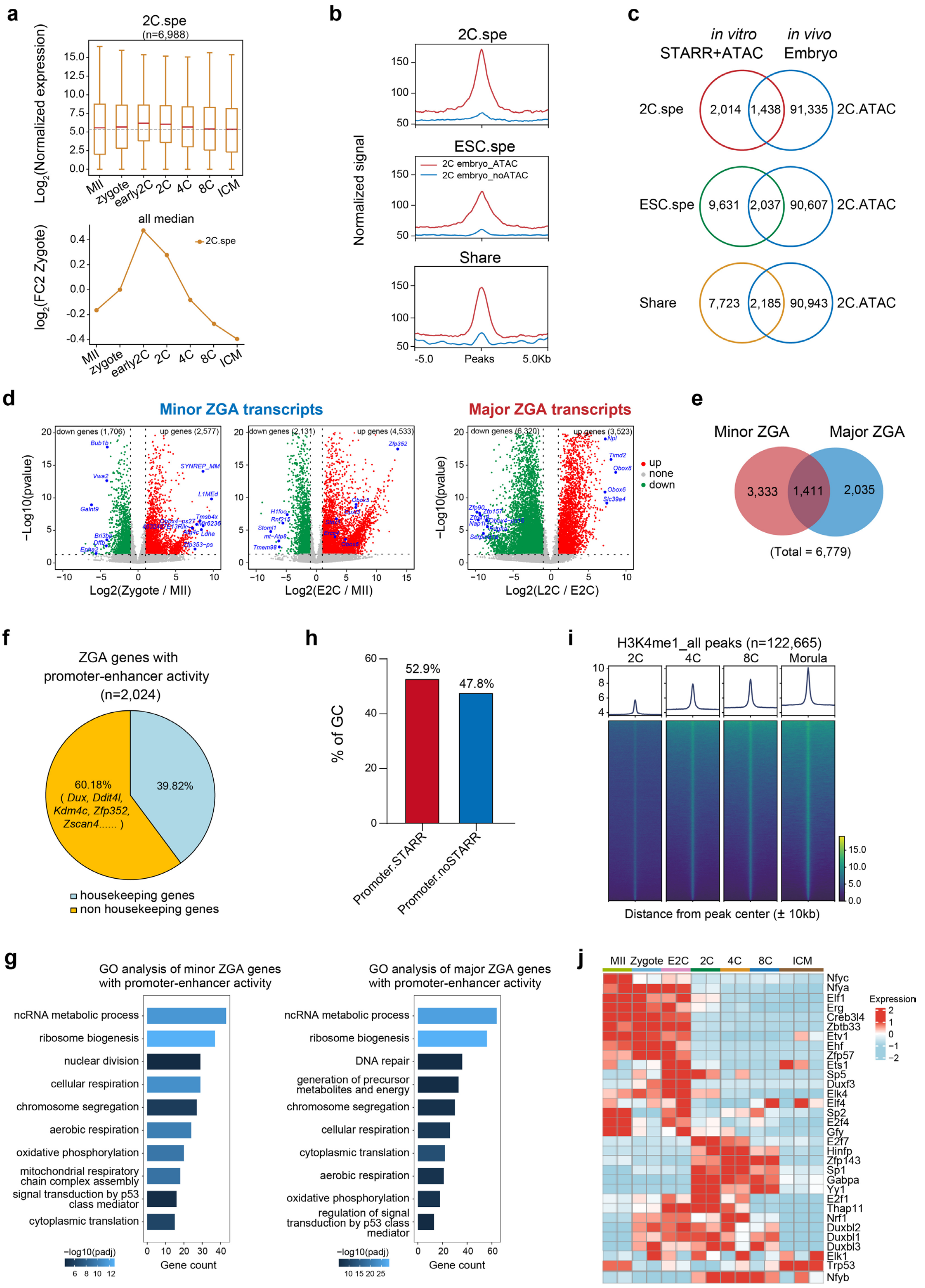

**Supplementary Fig. 4 | Evaluation of the *in vivo* functional potential of enhancers identified in 2CLCs. Related to Fig. 2.**

**(a)** Boxplot showing the expression levels of the genes flanked by indicated groups of STARR peaks within 20 kb (± 20kb) in MII oocytes and mouse preimplantation embryos at various developmental stages (top). Line chart showing the ratios of gene expression level relative to zygote at various embryonic stages (bottom). Line chart represents the medians as shown in the above Boxplot. The associated gene numbers were indicated. The RNA-seq data were obtained from published data (GSE66390).

**(b)** Line chart showing the ATAC-seq signals around 2C-specific (top), ESC-specific (middle) or shared STARR peaks (bottom) in mouse 2C embryos, categorized by the high (ATAC) or low (noATAC) ATAC-seq signals in 2C embryos. Signal was computed on the STARR-seq peaks flanked by 5 kb.

**(c)** Veen diagram showing the overlapping between ATAC peaks in mouse 2C embryos and the indicated clusters of STARR peaks. 2C.spe: 2CLC-specific STARR peaks (top); ESC.spe: ESC-specific STARR peaks (middle); Share: 2CLC/ESC shared STARR peaks (bottom).

**(d)** Volcano plot showing the differential expressed transcripts, including annotated genes and transposable elements (TEs), between zygote and MII oocyte (left), early 2-cell (E2C) and MII oocyte (middle), as well as late 2-cell (L2C) and E2C (right). The annotated genes upregulated in zygote or E2C compared to MII oocyte were regarded as minor ZGA genes, and the annotated genes upregulated in L2C compared to E2C were regarded as major ZGA genes. The plots include thresholds for significance, with a fold change greater than 1 and a p-value less than 0.01 indicated. The RNA-seq data were derived from published data (GSE66390).

**(e)** Veen diagram showing overlapping between minor (red, n = 4,744) and major ZGA genes (blue, n = 3,446) from Supplementary Fig. 4d. The total number of ZGA genes was 6,779.

**(f)** The proportion of housekeeping and non-housekeeping genes in the ZGA genes with 2CLC STARR-seq signals (n = 2,024) at promoter regions (TSS ± 1kb).

**(g)** Gene ontology (GO) enrichment analysis of minor (left) or major (right) ZGA genes with 2CLC STARR-seq signals at promoter regions (TSS ± 1kb).

**(h)** The GC content of the promoter sequences (Promoter.STARR and Promoter.noSTARR). Promoter.STARR: the ZGA gene promoters with STARR-seq signals in 2CLCs; Promoter.noSTARR: the ZGA gene promoters without STARR-seq signals in 2CLCs.

**(i)** Heatmap of global H3K4me1 levels in 2C, 4C, 8C and morula embryos. Signal was computed on the H3K4me1 peaks flanked by 10 kb.

**(j)** The RNA expression level of the TFs (from Fig. 2g) during mouse embryonic development. The TFs are from Fig. 2g, with their motifs enriched at ZGA gene promoters (TSS ± 1kb) with STARR-seq signals in 2CLCs.

#### Supplementary Fig. 5

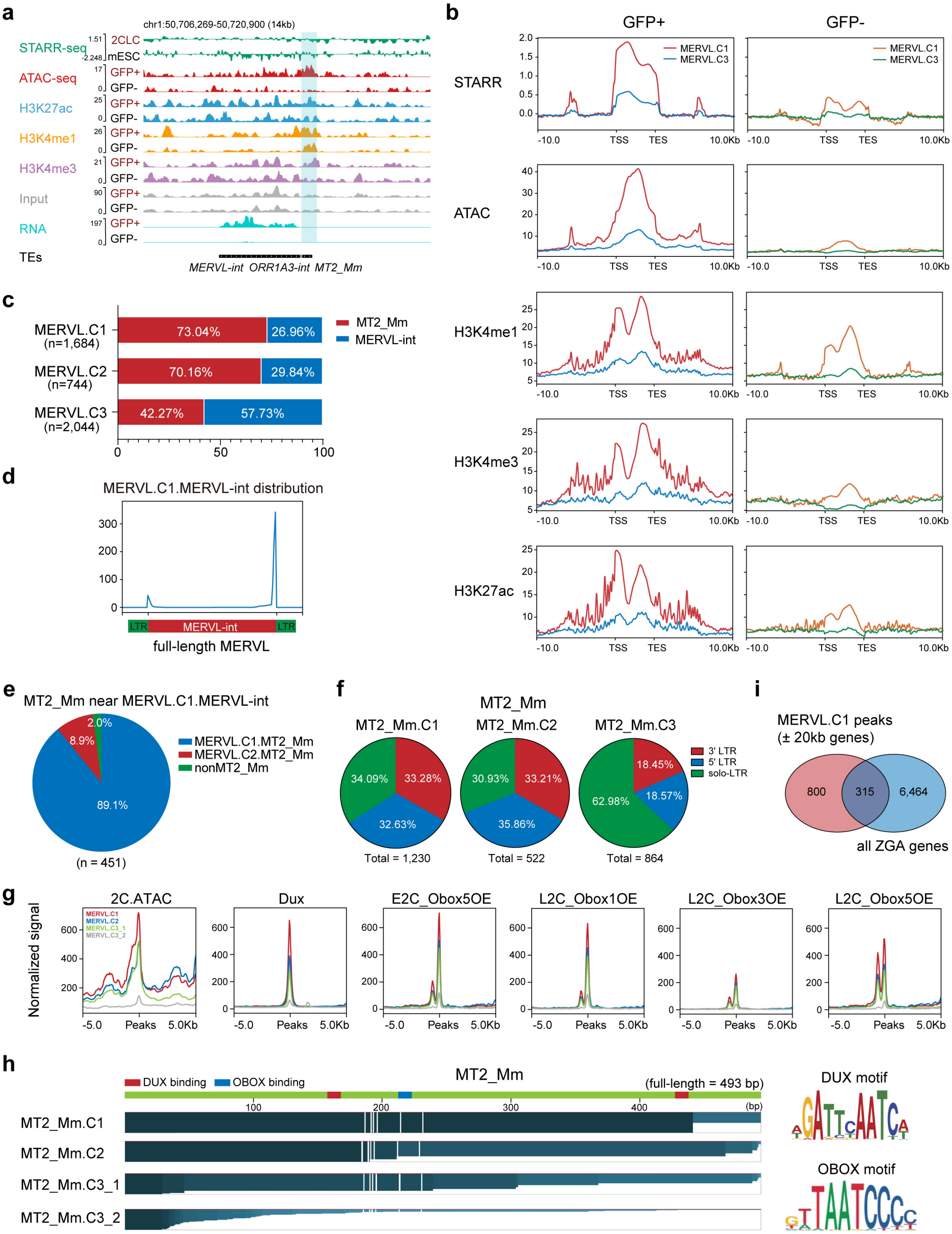

**Supplementary Fig. 5 | Features of MERVL elements with enhancer activities. Related to Fig. 3.**

**(a)** Genomic snapshot showing STARR-seq, ATAC-seq, ChIP-seq of the specified histone modification with input, and RNA-seq tracks at the indicated transposable elements (TEs).

**(b)** Line charts showing STARR, ATAC, H3K27ac, H3K4me1 and H3K4me3 signals around the indicated clusters of MERVL elements (± 10 kb) in MERVL-GFP$^{+}$ (left) and MERVL-GFP$^{-}$ cells (right).

**(c)** The distribution ratio of MT2_Mm and MERVL-int in the indicated three clusters of MERVL elements in Fig. 3d.

**(d)** The location distribution of MERVL-int fragments within MERVL.C1 in the full-length MERVL sequence.

**(e)** The proportion of MERVL-int fragments within MERVL.C1 located near the indicated clusters of MT2_Mm.

**(f)** Pie charts showing the proportion of three different categories of MT2_Mm in the indicated clusters of MERVL elements in Fig. 3d.

**(g)** Line charts showing the ATAC-seq signals and ChIP-seq signals of different TF binding at the indicated clusters of MT2_Mm (Fig. 3h) in mouse 2C embryos. E2C: early 2C embryos; L2C: late 2C embryos.

**(h)** The fragment length of the indicated four clusters of MT2_Mm elements in Fig. 3h (left). The potential binding sites of DUX (red) and OBOX (blue) on MT2_Mm are marked in the schematic diagram. The DUX and OBOX motifs were shown (right).

**(i)** Venn diagram showing overlapping between all ZGA genes and the genes flanked by MERVL.C1 peaks within 20kb (± 20kb).

#### Supplementary Fig. 6

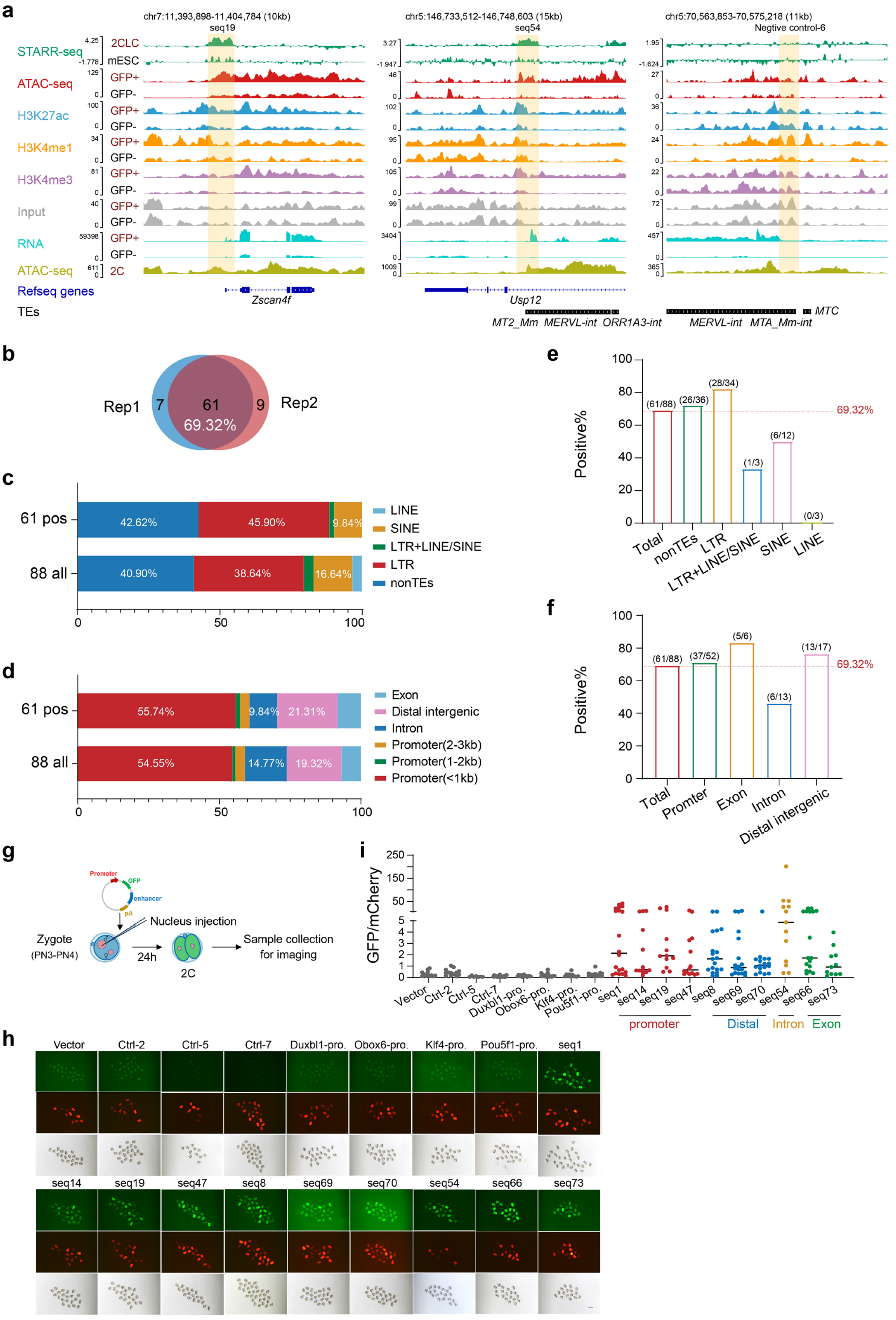

**Supplementary Fig. 6 | Verification of the 2CLC enhancers in 2C embryos. Related to Fig. 4.**

**(a)** Genomic snapshots showing STARR-seq, ATAC-seq, ChIP-seq of the specified histone modification with input, and RNA-seq of indicated histone modification and RNA-seq tracks at the indicated genome loci. The highlighted regions marked the representative enhancer candidates (seq19 and seq54) and negative control which were selected for ULI-STARR-seq in 2C embryos.

**(b)** Venn diagram showing the number of overlapping positively enriched enhancer candidates between the two replicates in Fig. 4b.

**(c)** The proportion of different TE classes in 88 enhancer candidates (all) and 61 positive enhancers (pos). LTR indicated the LTRs of transposable elements.

**(d)** The genomic distribution of 88 enhancer candidates (all) and 61 positive enhancers (pos).

**(e)** The percentage of positive enhancers within the specified groups of enhancer candidates, categorized by their associated transposable elements (TE) as depicted in Supplementary Fig. 6c.

**(f)** The percentage of positive enhancers within the specified groups of enhancer candidates, categorized by their genomic distribution as depicted in Supplementary Fig. 6d.

**(g)** Schematic illustration of pronuclear injection with the enhancer reporter assay. The embryos were collected for fluorescence analysis at 24 h post-injection.

**(h)** Fluorescence and bright fields of mouse embryos in the enhancer reporter assay. Pro.: Promoter. Scale bar: 100 μm.

**(i)** Dot plot showing the ratio of GFP to mCherry intensity in the enhancer reporter assay. The numbers of embryos used in each group: 12, 13, 7, 20, 20, 13, 14, 11, 14, 20, 13, 12,13, 14, 17,19, 18, 17, 13, 17 and 11.

**Supplementary Fig. 7**

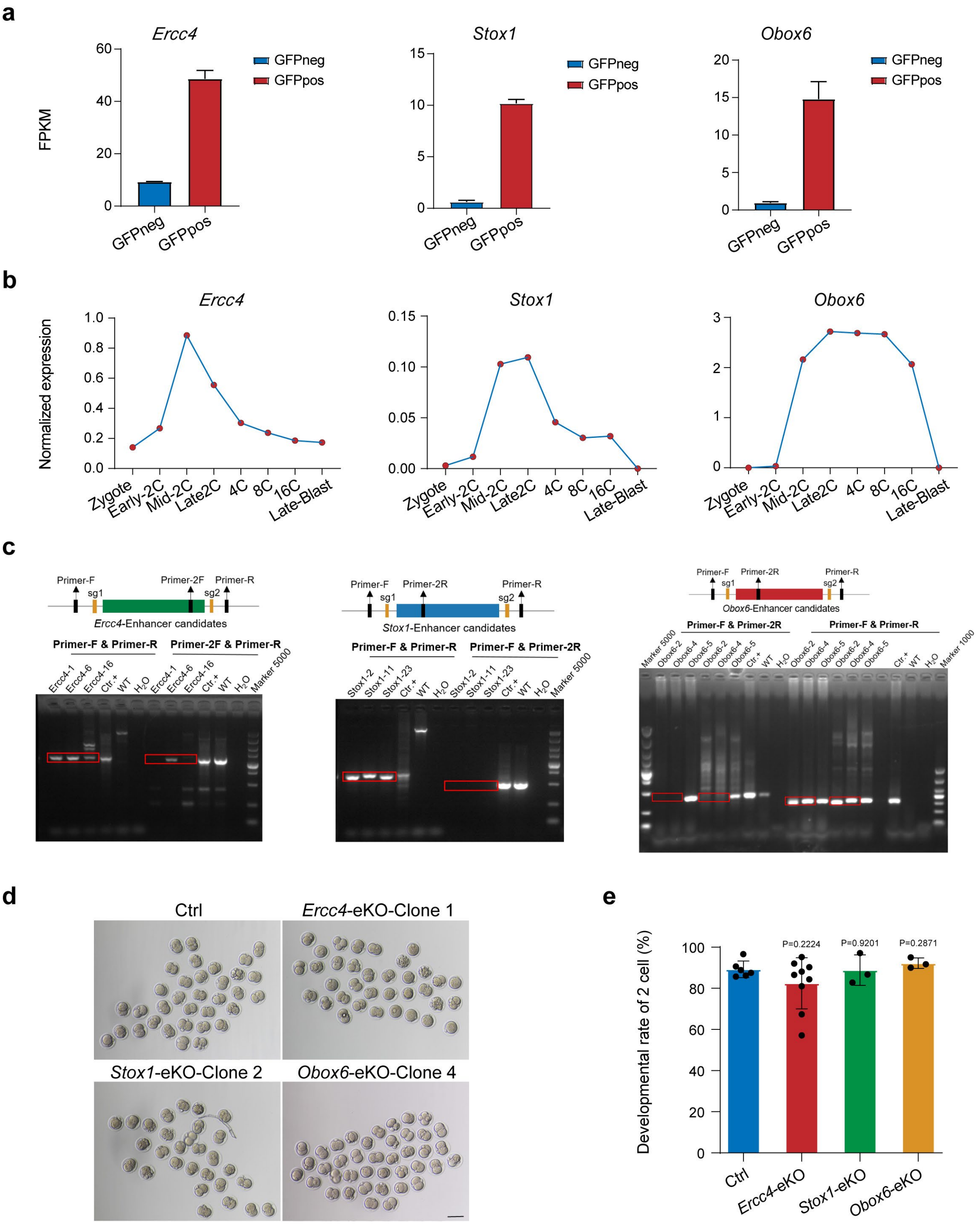

**Supplementary Fig. 7 | Verification of the enhancer functionality by CRISPR/Cas9 knockout. Related to Fig. 4.**

**(a)** Boxplot showing the RNA expression levels of three indicated genes (also seen Fig. 4d) in MERVL-GFP$^-$ and MERVL-GFP$^+$ cells. Values are means ± SD, n = 3 biological replicates.

**(b)** Line chart showing the changes in RNA expression levels of three indicated genes during mouse embryonic development. Published RNA-seq data were used (GSE66390).

**(c)** Genotyping strategies (top) and gel pictures (bottom) of the genotyping of indicated samples including single cell-derived eKO ESC clones. eKO: enhancer knockout. Ctr.+: positive controls with bulk-population cells transfected with Cas9 and corresponding small gRNAs (sg); WT: wildtype cells transfected with Cas9 only; $H_2O$: water.

**(d)** Representative images of 2-cell stage control and eKO embryos generated by nuclear transfer with control cells (cells transfected with Cas9 only) and specified enhancer knockout (eKO) clones. Red arrows indicate the embryos arrested at 1-cell stage. Scale bar: 100 µm.

**(e)** The developmental rate of late 2C embryos is shown for both the control and indicated eKO groups, normalized to the control group within each replicate. eKO: enhancer knockout. Values are means ± SD, n ≥ 3 biological replicates. Two-sided paired t tests were performed.
